# Genetic basis of an adaptive polymorphism controlling butterfly silver iridescence

**DOI:** 10.1101/2024.12.13.628425

**Authors:** Luca Livraghi, Joseph J. Hanly, Ling Sheng Loh, Albie Henry, Chloe M.T. Keck, Vaughn M. Shirey, Cheng-Chia Tsai, Nanfang Yu, Steven M. Van Belleghem, W. Mark Roberts, Carol L. Boggs, Arnaud Martin

## Abstract

Identifying the genes and mutations that drive phenotypic variation and which are subject to selection is crucial for understanding evolutionary processes. Mormon Fritillary butterflies (*Speyeria mormonia*) exhibit a striking wing color polymorphism throughout their range: typical morphs bear silver spots on their ventral surfaces, and can co-occur with unsilvered morphs displaying a dull coloration^1^. Through genome-wide association studies in two polymorphic populations, we fine-map this difference in silvering to the 3’ region of the transcription factor gene *optix*. The expression of *optix* is confined to the unsilvered regions that surround the spots, and these patterns are transformed to a silver identity upon *optix* RNAi knockdown, implicating *optix* as a repressor of silver scales in this butterfly. We show that the unsilvered *optix* haplotype shows signatures of recent selective sweeps, and that this allele is shared with the monomorphic, unsilvered species *Speyeria hydaspe*, suggesting that introgressions facilitate the exchange of variants of adaptive potential across species. Remarkably, these findings parallel the role of introgressions and *cis*-regulatory modulation of *optix* in shaping the aposematic red patterns of *Heliconius* butterflies^2–7^, a lineage that separated from *Speyeria* 45 million years ago^8^. The genetic basis of adaptive variation can thus be more predictable than often presumed, even for traits that appear divergent across large evolutionary distances.

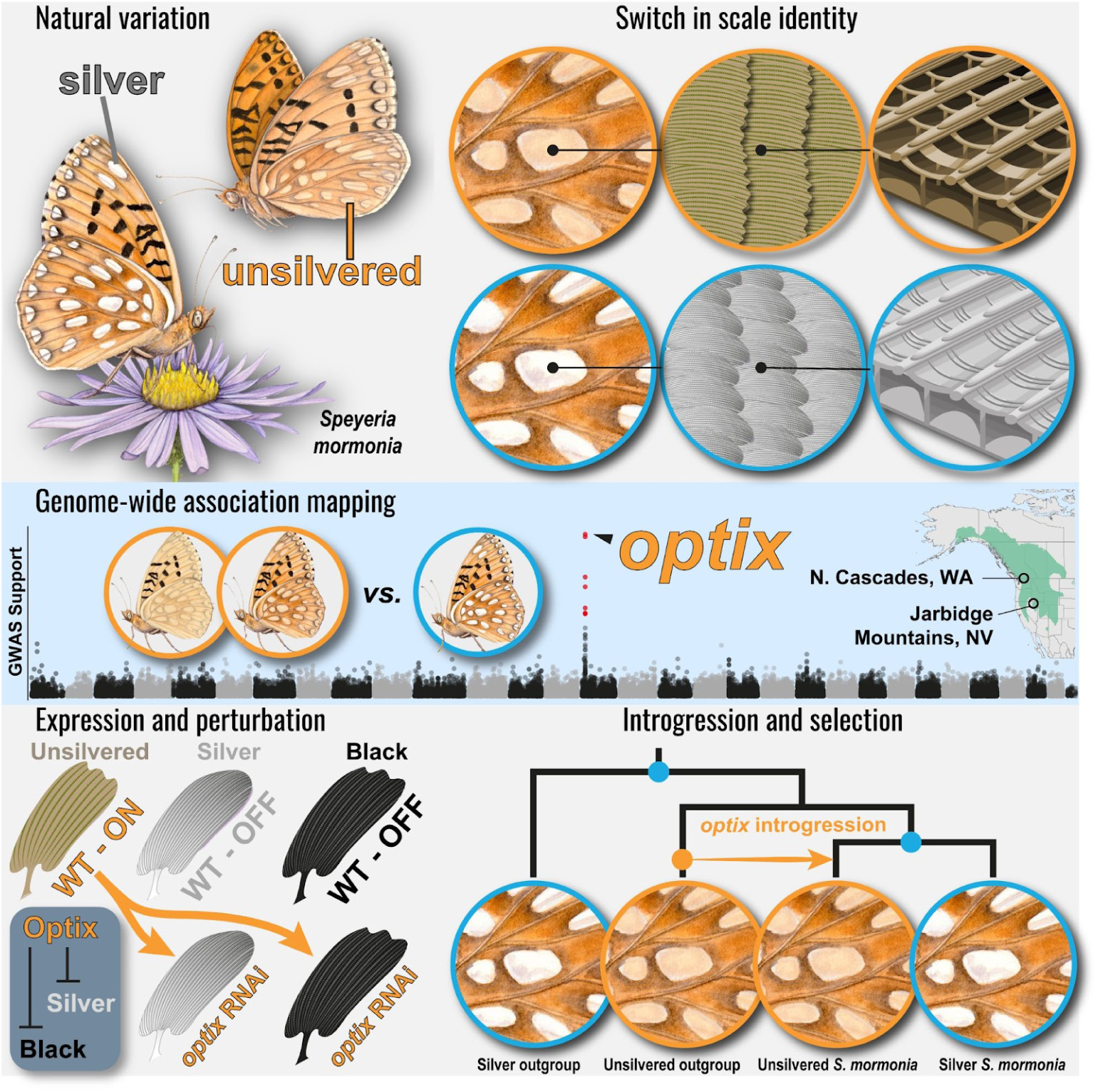
Graphical Abstract.

## Results

### Complex spatial distribution of silver polymorphism in *Speyeria mormonia*

The ventral hindwings of *Speyeria (Argynnis) mormonia* display a polymorphism for the presence of iridescent silver scales^9^ (**Figures 1A and 1A’**). Unsilvered morphs have spots with a light beige color, while the same patterns have a bright metallic look in silver morphs (**Figures 1B-1B’ and S1**). The difference between silver and unsilvered spots is due to a scale identity shift (**Figures 1C-1D’**): most notably, unsilvered scales are slightly pigmented and porous (light-absorbent), while silver scales are unpigmented and with a continuous upper surface (light-reflective)^10–12^. *Speyeria mormonia* occurs across mountainous habitats of western North America, with silver and unsilvered morphs varying in frequencies throughout its range^13^. However, despite several studies reporting similar polymorphisms in many species of *Speyeria*^14,15^, none thus far have systematically described their geographic distribution. To accurately map the geographic variation of the silver scale polymorphism in *S. mormonia*, we examined 9,760 records from museum collections across the US and Canada, scoring the prevalence of silver and unsilvered morphs from each locality. We observed a complex, heterogeneous landscape of phenotype distributions, with a general decrease in silver morph frequency with increasing latitude (**Figure 1E and Figure S2**). Local heterogeneity in unsilvered/silver morph frequency is apparent throughout the species range. For example, unsilvered morphs (*S. mormonia artonis*) are prevalent in populations from the Steens, Jarbidge and Ruby Mountain ranges in southeastern Oregon and northern Nevada, in sharp contrast to surrounding localities where these morphs become rare (**Figure S3**). Regression analyses between these trait proportions and geoclimatic variables found weak associations (R^2^ = 0.32-0.33) with solar irradiation and isothermality (**Figure S2**), two variables that are directly determined by latitude. Yet, mid-infrared imaging shows that silver spots do not increase the thermal emissivity of these wing patterns (**Figure S4**), ruling out a radiative cooling effect^16^ that may have explained their higher prevalence at lower latitudes. Because none of the tested environmental variables were a strong predictor of observed silvering proportions, we conclude that the observed heterogeneity in prevalence of each morph is influenced by local demographic factors, or by micro-habitat differences that do not scale to the larger distribution range of the species.

**Figure 1.**
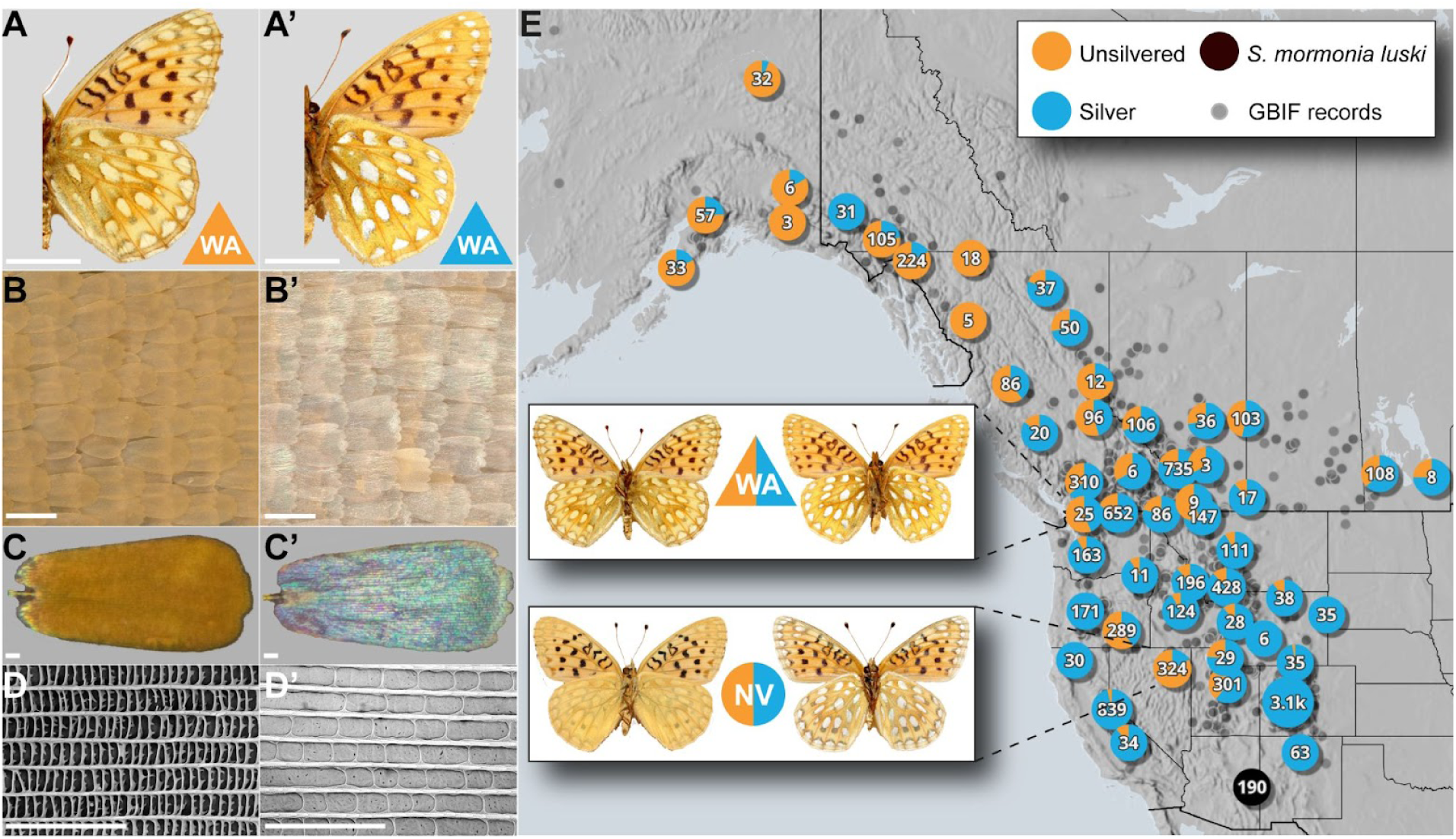
Ultrastructural basis and geographic distribution of the silvering polymorphism in *Speyeria mormonia*. (**A-A’**) Representative *S. mormonia* individuals exhibiting unsilvered and silver ventral phenotypes. **(B-B’)** Magnified view of unsilvered and silver wing spots. **(C-C’)** Reflected light micrographs of single scales from each morph; unsilvered scales are brown, while silver scales are unpigmented but iridescent. **(D-D’)** Scanning electron micrographs of the upper surfaces of unsilvered and silver scales. (D) Unsilvered scales have a perforated upper lamina that makes them non-reflective, while (D’) silver scales have a continuous upper lamina that makes them reflective. **(E)** Geographic distribution of the silvering polymorphism. Pie charts indicate the observed proportions of silver (blue) and unsilvered (orange) individuals and report the total number of scored individuals at each location. A distinct subspecies, *S. mormonia luski*, found in Arizona and exhibiting an atypical white-spotted phenotype, is indicated in black. The polymorphic populations sampled in this study for sequencing are highlighted: one from the North Cascades in Washington (WA) and another from the Jarbidge Mountains in Nevada (NV), with representative images of silver and unsilvered individuals shown. Gray points represent 9,717 occurrence records of *S. mormonia* from GBIF that lack phenotypic information, or for which silvering was not scorable from pictures. Scale bars: A-A’ = 1 cm; B-B’ = 100 μm; C-D’= 10 μm.

### The silvering polymorphism maps to *optix*

To determine the inheritance patterns and dominance hierarchy of silvering, we crossed captive-reared *S. mormonia* individuals from a mixed population in Manning Park, British Columbia, generating 931 individuals from 26 broods over 3 generations. Offspring phenotypic frequencies indicate that silver iridescence is inherited as a single-locus, recessive autosomal trait (**Table S1**), consistent with observations in other *Speyeria* species^14,15^.

Given the simple genetic basis for the silver/unsilvered switch, we sought to map the underlying locus by Genome-Wide Association (GWA). To achieve this, we resequenced the genomes of 70 individuals sampled from independent admixed populations in the North Cascades (WA) and in the Jarbidge Mountains (NV) – where silver individuals are found at around 40% frequency (**Figure S2**). Following single nucleotide polymorphism (SNP) calling against an existing genome assembly of *S. mormonia*^8^, the GWA study identified a ∼7-kb interval on autosome 14 strongly associated with the presence/absence of silver scales (**Figure 2A**). The associated SNPs map to an intergenic region ∼15 kb downstream of the transcription start site of *optix,* a gene with a well-documented and conserved role in butterfly scale pigmentation^3,17–19^ (**Figure 2B**). The statistical support for the GWA SNPs at the *optix* locus increases in the combined NV and WA datasets when population structure is accounted for (**Figure S5A-B**). A secondary GWA signal found exclusively in the NV population maps to a *DIRS1-like* transposable element insertion site on chromosome 25 (**Figures 2A and S5C**). Due to the repetitive nature of this 5-kb region (11 copies in the reference genome) and the limitations of short-read genotyping, we cannot confirm the validity of this mapping result and excluded it from further analyses.

**Figure 2.**
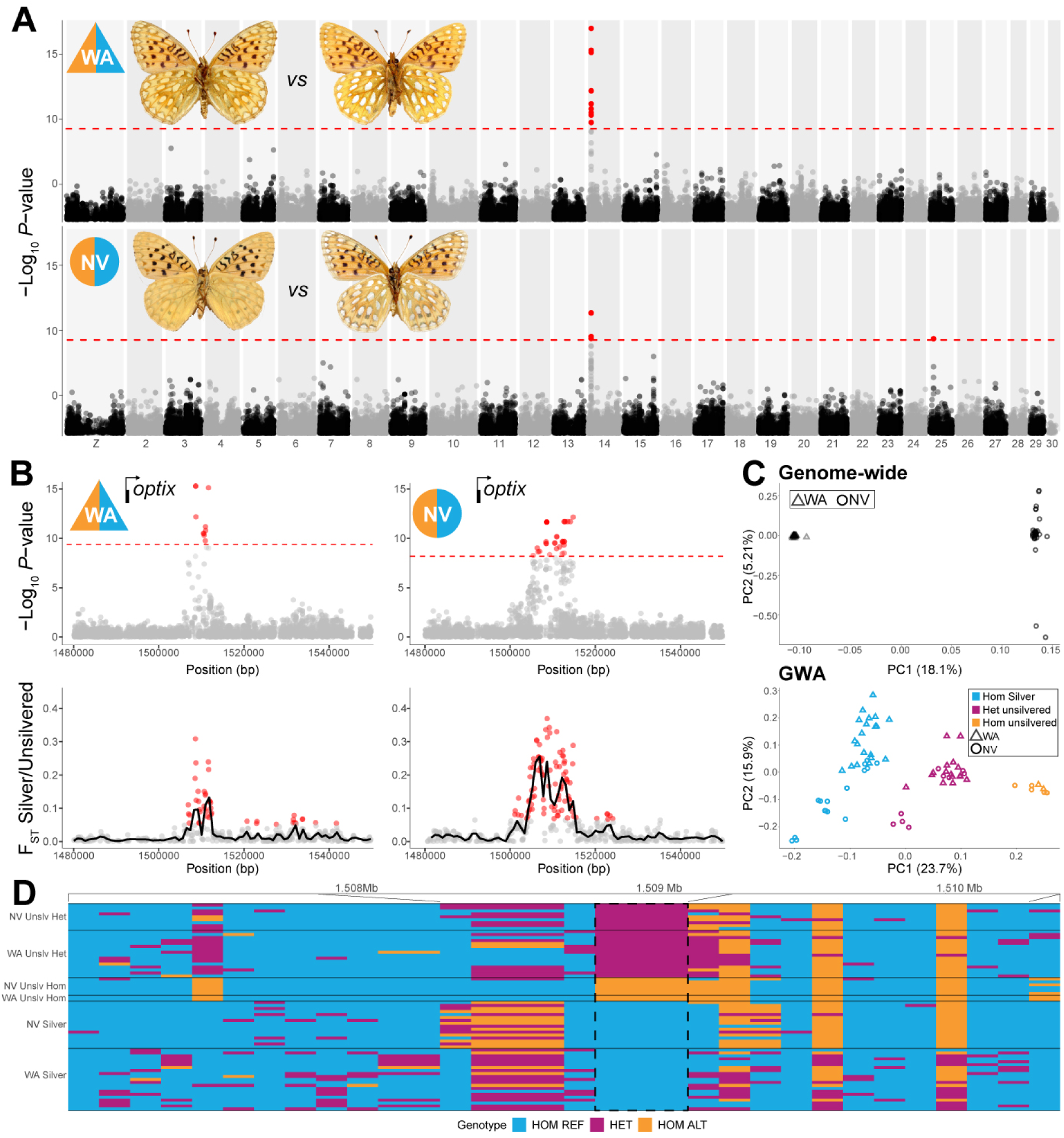
The silvering locus in *S. mormonia* maps to the *optix* transcription factor gene. **(A)** GWAS of the silvering polymorphism reveals a strong association on chromosome 14 in both the WA and NV populations. Chromosome numbers are indicated at the bottom. The y-axis shows the Wald test negative log_10_ *P*-value associated with each variant. The dotted line represents the Bonferroni corrected genome-wide significance threshold (*p* < 0.01). **(B)** The associated SNPs are located ∼15 kb downstream of the *optix* transcription start site (black arrow), and are accompanied by moderate differentiation between silver and unsilvered individuals as measured by F_ST_, calculated in 250-bp sliding windows. F_ST_ windows in the 99th percentile of the chromosome-wide distribution are shown in red. **(C)** Principal component analysis (PCA) based on genome-wide SNPs clusters samples by geography (PC1). Performing a PCA using the GWA interval clusters the samples by phenotype along PC1, and by geography on PC2. **(D)** Genotype plot for the GWA region. Each row is an individual, and each column is a color-coded SNP. Three fixed SNPs are found between silver and unsilvered individuals in both populations (highlighted by black dashed box).

Next, we more closely examined patterns of SNP sharing within the chromosome 14 GWA interval in each sampled population. The association signal at *optix* is accompanied by an increased but not fixed F_ST_ signal, driven by high frequencies of heterozygous unsilvered individuals (**Figures 2C-2D**). Genome-wide principal component analysis (PCA) largely clusters individuals by geography, whereas local PCA in the GWA interval clusters individuals by phenotype along PC1, with PC2 recovering some geographic signal (**Figure 2C**). Scoring SNPs by zygosity state within the GWA interval identified 3 SNPs differing between unsilvered and silver butterflies, with the majority of unsilvered *S. mormonia* being heterozygous for these fixed SNPs (**Figures 2D and S6**).

### *Optix* represses silver scale specification

To test the developmental role of *optix* in silver pattern formation, we reared silver individuals collected from Colorado, a location where silver alleles are nearly fixed^20^, and examined *optix* expression by hybridization chain reaction (HCR). In early pupal wings (28% stage), *optix* expression is restricted to two small patches of presumptive “coupling scales” (**Figure 3A**) – the future zone of contact between forewings and hindwings – consistent with previous studies in other butterflies^11,19^. At 54% of pupal development (**Figure 3B**), *optix* shows widespread expression in developing scales but is markedly absent from the presumptive silver hindwing spots, as well as from various presumptive black pattern elements. These results suggest that *optix* is repressed in patterns with either silver or melanic scales, and may thus act as a repressor of these differentiated states in *S. mormonia*.

**Figure 3.**
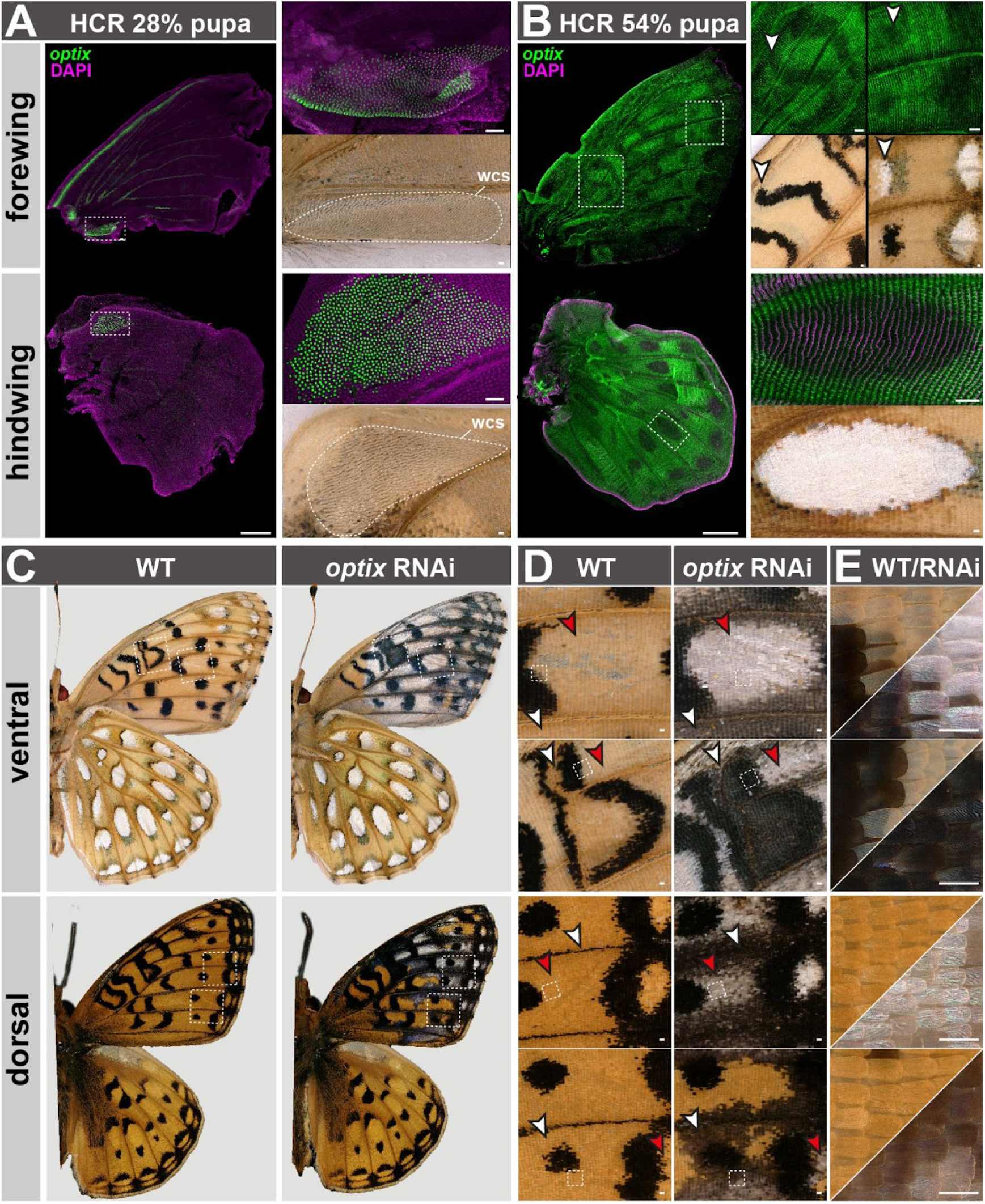
*Optix* represses silver and black scale specification. **(A)** HCR stainings in 28% pupal wings show *optix* expression in the presumptive wing coupling scales (wcs), a patch of tooth-shaped scales that create friction in a zone of contact between the forewing and hindwing. **(B)** By 54% of pupal development, *optix* is localized throughout the wing, except in the presumptive silver and black pattern elements (examples indicated with arrowheads). **(C-E)** Electroporated RNAi knockdowns of *optix* result in homeotic scale transformations from orange to silver (red arrowheads) or black (white arrowheads) on both wing surfaces. Wild-type (WT) and RNAi-treated images correspond to the left and right side to the same individuals, with the WT horizontally flipped to facilitate comparisons here. Additional RNAi knockdown replicates are shown in **Figure S7**. Scale bars: A-B, left panels = 1 mm; A-B (right panels) = 100 μm; D-E = 100 μm.

Next we sought to perturb *optix* to test this hypothesis. To achieve this, we used pupal wing electroporation of Dicer-Substrate Short Interfering RNAs (DsiRNAs)^21^ instead of CRISPR mutagenesis, bypassing the need for a breeding colony by allowing experiments in field-collected larvae that survived their obligatory diapause. Remarkably, RNA interference (RNAi) knockdowns of *optix* resulted in a shift from non-reflective scale states to silver, showing that *optix* is necessary to prevent silver scale specification (**Figure 3D-E**, red arrows). We also observed position-dependent effects, where orange areas transformed to black, rather than silver (white arrows), thus implying that *optix* combines with other unknown factors to determine color scale identity. In summary, *optix* is downregulated in silver spots and represses silver scale identity, confirming its function as a switch gene that causes the silvering polymorphism in natural populations.

### The unsilvered allele shows signatures of selective sweeps

The observed variation in polymorphism frequencies across populations hints at the influence of non-neutral evolutionary forces maintaining this genetic diversity. The recessive silver state is nearly fixed in the Cascade and Klamath Mountain ranges (OR, North CA) and in the Eastern Rockies (CO, NM, SD). In contrast, the populations we sampled in WA and NV show intermediate and high frequencies of the dominant unsilvered allele (**Figure S3**). To better understand these dynamics, we carried out differentiation and selection scans of the *optix* silvering locus. Using the three fixed SNPs identified in the GWAS as diagnostic of either unsilvered or silver haplotypes (**Figure 2D**), we generated a set of phased SNPs on chromosome 14 and coded each haplotype as associated to unsilvered or silver states. Recalculating F_ST_ in sliding windows on these phased haplotypes shows an elevated signal, reflective of the higher differentiation between individual alleles (**Figures 4A and 4A’**). Allele divergence is also higher at the *optix* locus, as shown by elevated Dxy (**Figures 4B and 4B’**), and nucleotide diversity (π) is reduced within the elevated F_ST_ window for unsilvered haplotypes in both populations (**Figures 4C and 4C’**). Lastly, Tajima’s D is also reduced over this genomic region for the unsilvered haplotypes only, indicating the presence of sites deviating from neutral evolution (**Figures 4D and 4D’**).

**Figure 4.**
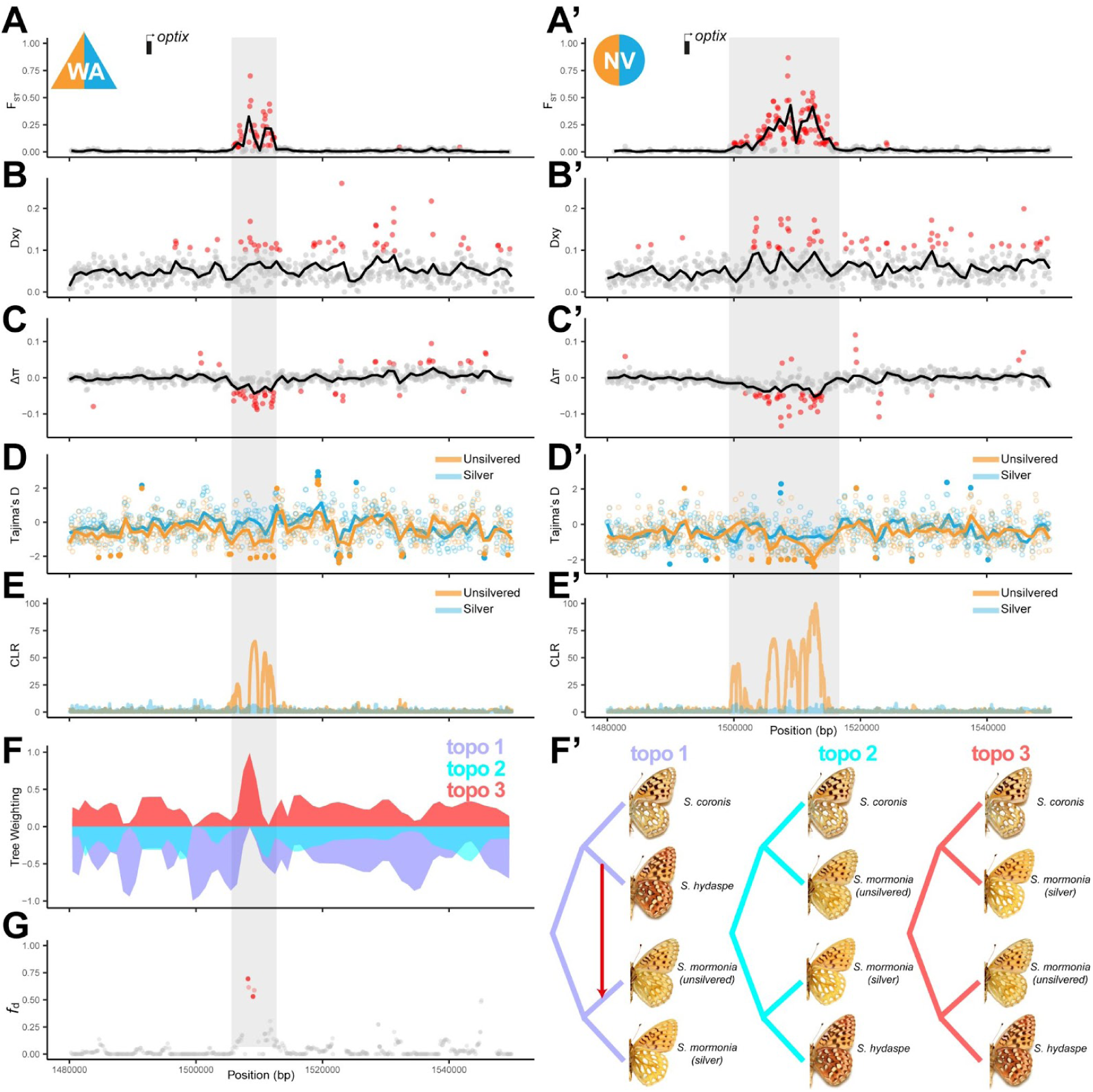
Evidence of selection and introgression at the unsilvered allele. (**A-A’**) Strong differentiation as measured by F_ST_ between individually phased unsilvered and silver alleles measured in 200-bp windows. This signal is observed at the 3’ end of *optix* in both WA (A) and NV (A’) populations. **(B-B’)** The same genomic interval is accompanied by locally elevated Dxy, and **(C-C’)** a reduction in nucleotide diversity (Δπ=π_unsilvered_ – π_silver_) whereby unsilvered alleles show a reduction in diversity compared to silver alleles. **(D-D’)** A reduction in Tajima’s D can also be observed for the buff haplotype concurrent with the F_ST_ and Δπ signals, indicating the presence of non-neutrally evolving sites. **(E-E’)** Haplotype-based selection statistics show elevated CLR at the locus for unsilvered alleles only. **(F-F’)** Phylogenetic weighting of phenotypes show strong support for shared alleles between *S. hydaspe* and unsilvered *S. mormonia* as measured by TWISST^25^. Topo1: expected species-level topology with monophyletic *S. mormonia*^27^; Topo3: topology grouping lineages by silvering state. **(G)** Proportion of introgression or gene flow between *S. hydaspe* and unsilvered *S. mormonia* (red arrow in F’), as measured by *f*_d_ shows an excess of shared sites between them, as compared to the silver outgroup *S. coronis*^27^. Red dots (A-C’, G) and filled dots (D-D’) highlight absolute values in the 99th percentile of their chromosome-wide distribution.

We next scanned the *optix* locus for selective sweeps using the composite-likelihood ratio (CLR) statistics implemented in SweeD^22^. These scans show selective sweeps at the unsilvered allele in both the WA and NV polymorphic populations (**Figures 4E and 4E’**), matching the position of the differentiated haplotypes, which span the 3 GWA SNPs and blocks of hitchhiking variants around them (**Figure S6**). No signatures of selective sweeps were found at the silver haplotypes. Because selection scans can only detect recent episodes of positive selection^23,24^, silver alleles may have been fixed at these locations in a more distant past, and thus accumulated genetic diversity that would obscure a potential ancient sweep signal.

### Adaptive introgression of the unsilvered allele

The genus *Speyeria* is relatively large and rapidly radiating, with several occurrences of sympatric species harboring monomorphic populations with either unsilvered or silver scales. In much of its range, *S. mormonia* occurs in sympatry with the closely related, monomorphic unsilvered species *Speyeria hydaspe*, and interspecies crosses in laboratory conditions between the two can produce fertile offspring^14,15^. We therefore asked whether occasional hybridization could result in introgressed unsilvered alleles into *S. mormonia* populations from *S. hydaspe*. To test for allelic sharing between the two species, we applied a topology weighting test (TWISST^25^), with 4 defined groups all sampled from the same WA North Cascade location: a silver outgroup (*Speyeria coronis*), unsilvered *S. hydaspe*, and unsilvered and silver phased alleles of *S. mormonia*. TWISST scans group the samples by phenotypic state rather than by species phylogeny at the GWA interval (**Figures 4F and 4F’**), suggesting identity by descent of the unsilvered *S. hydaspe* and *S. mormonia* alleles. We then measured the proportion of introgression or gene flow between unsilvered *S. mormonia* and *S. hydaspe* by calculating the *f*_d_ statistic in sliding windows^26^ and found support for introgression at sites concurrent with elevated F_ST_ and CLR (**Figure 4G**). These results suggest that the unsilvered alleles of *S. mormonia* originated by adaptive introgression from an unsilvered, diverged species.

## Discussion

### Genetic repeatability of adaptation in diverged traits

Is the genetic basis of adaptation predictable^28,29^? Empirical studies of phenotypic convergence suggest it is to some extent, as the same genes are often found to drive the repeated evolution of traits with simple genetic architecture, or show parallel signatures of differentiation across populations that have adapted to similar ecological challenges^30,31^. This evolutionary repeatability or “gene reuse”^30^ can arise through independent mutations at the same hotspot loci, the introgression of adapted alleles across diverged gene pools, or the incomplete lineage sorting of standing genetic variation (ILS). Here, we found that the *optix* locus drives adaptive color pattern variation at levels of replication that span the micro and macroevolutionary scales, depicting a nuanced view of how the genotype-phenotype map shapes phenotypic diversity.

At the microevolutionary level, unsilvered alleles are identical across two distinct *S. mormonia* populations, and introgressed from an unsilvered species of the same genus. At the macroevolutionary level, our findings in *S. mormonia* parallel the repeated involvement of *optix* alleles in shaping adaptive variation in *Heliconius* and *Hypolimnas*^2,3,24,32–34^, two lineages that diverged form *S. mormonia* 45-70 MYA^8,35^. In these two other lineages, *optix* variation does not control silvering and camouflage strategies, but instead switches the phenotypic expression of black to red and orange color patterns used in toxic mimicry. It is noteworthy that the *optix* locus has already revealed multiple cases of adaptive introgression within the *Heliconius* radiation, as well as evidence that independently evolved *cis*-regulatory modules recombined to make new red patterns^4,7,36,37^. We have now found a second lineage where *optix* alleles can introgress between diverged species and then be maintained by selection. Thus, patterns of silvering diversity within New World species of the *Speyeria* genus, within and between species, may be explained by the repeated selection of the alleles that exist as ancient standing genetic variation (ILS), or via hybridization between inter-fertile lineages. Future work will be required to clarify the extent to which this variant is spread across other polymorphic *Speyeria* lineages^38^, including in related Old World species of the *Argynnis/Fabriciana* genus^39^.

Remarkably, the reuse of *optix* across macroevolutionary time invokes convergent genetics, but without phenotypic convergence: how does the same locus drive two distinct effects, red-orange pattern variation in *Heliconius/Hypolimnas*, and silver dimorphism in *Speyeria*? Elements of an answer come from perturbation studies that examined the phenotypic effects of *optix* loss of function with CRISPR knock-outs and RNAi^11,17,33,40^. In short, *optix* acts as a master selector gene that can switch both the pigmentary and ultrastructural identity of wing scales. First, all species examined so far show that *optix* is necessary for activating ommochrome pigmentation (red-orange scales) while also repressing dark melanin outputs in these same scales. In other words, *optix* deficiency results in a conversion of red-orange patterns to black in at least some parts of the wings^17,33,40^, including in *S. mormonia* (**Figures 3C and 3D**). This function of *optix* in upregulating ommochromes while repressing melanic states thus appears conserved, but varies in its spatial distribution across species. Second, *optix* also modulates scale ultrastructures, as it is required for the specification of wing coupling scales in several species^11,41^. In *Junonia coenia*, *optix* KOs also result in a gain of blue iridescence by thickening the scale’s lower lamina^41,42^. In *S. mormonia*, we found that WT silver scales lack *optix*, and that buff and orange scales convert to reflective silver upon *optix* knockdown, implying an increase in upper lamina coverage that explains their high reflectance (the reverse from its effect in wing coupling scales). The collective evidence shows that *optix* controls the deposition of chitin in the lower or upper laminae, resulting in changes in scale reflectance and iridescence, in both taxon– and position-dependent manners.

These data establish the *optix* transcription factor gene as a versatile genetic tool for the evolutionary diversification of wing patterns, including distinct adaptive polymorphisms involved in camouflage and mimicry across three lineages separated by 45-70 MY of divergence. More broadly, selector genes like *optix* are thought to act as regulatory hubs that integrate positional information and orchestrate the regulation of cell phenotypes. These so-called “input-output” genes may be privileged mutational targets of morphological evolution by combining evolvable expression with coordinated downstream effects on cell identity^29^. We extrapolate that similar genes can drive the repeated evolution of both convergent and divergent phenotypes, by combining a *cis*-regulatory architecture that facilitates their spatial refinement (as seen in the diverse expression patterns of *optix* across butterflies), and an inherent capacity to switch cell responses in a given tissue (*e.g.*, dual roles in pigmentary and ultrastructural outputs).

### Limitations of the study

While we could not procure the developmental samples to test this, we predict that unsilvered morphs express *optix* in the unsilvered spots. Testing this would clarify if GWA SNPs control *optix* recruitment in the spots in a spatial or temporal manner. Another expectation from this working model is that RNAi knockdowns should have strictly equivalent effects between the two morphs.

Importantly, the ecological functions of the silver and unsilvered wing spots remain to be formally studied. A role in thermoregulation is unlikely, because *S. mormonia* silver spots do not show noticeable heat-dissipating properties (**Figure S4**). Wing spots may be involved in intraspecific signaling (courtship, male competition, or species recognition), but a differential effect of the two forms may be difficult to detect. The exposure of the polymorphic pattern at the tip of the forewing (*i.e.* following Oudemans’ principle^43^) suggests a function when the butterfly is at rest (**Figure S1**). Unsilvered morphs are likely cryptic, with beige and brown features potentially matching visual backgrounds of dirt and dry vegetation (**Figure S1C**). Silver spots, while more conspicuous, might make edge recognition difficult and represent a form of disruptive camouflage against vertebrate predators^44,45^, or might be used to deter or confuse invertebrate predators^46,47^.

## Acknowledgements

We are grateful to Crispin Guppy, Norbert Kondla, Paul Hammond, Jonathan Pelham, Maru Losada, Randy Mooi, and Emma Eakins for providing invaluable collection records and phenotypic data, as well as the curation staff of several entomology collections and many iNaturalist and Butterfly eSurveys online contributors and volunteers (as listed in **Table S1**). We thank Kathleen Reinhardt, Erica Fleishman and Blair Stokes for assistance with specimen collection; the GWU HPC team for providing computing infrastructure; the GW Nanofabrication and Imaging Center (GWNIC), Patricia Hernandez and Aleksandar Jeremic for microscopy resources. Watercolor paintings featured in the Graphical Abstract were made by Julie Johnson (Life Science Studios). This work was supported by the NSF award IOS-2110534 to AM, a GWU University Facilitating Fund seed grant, and an EECG Embarkation grant from the American Genetics Association to LL.

## STAR★Methods

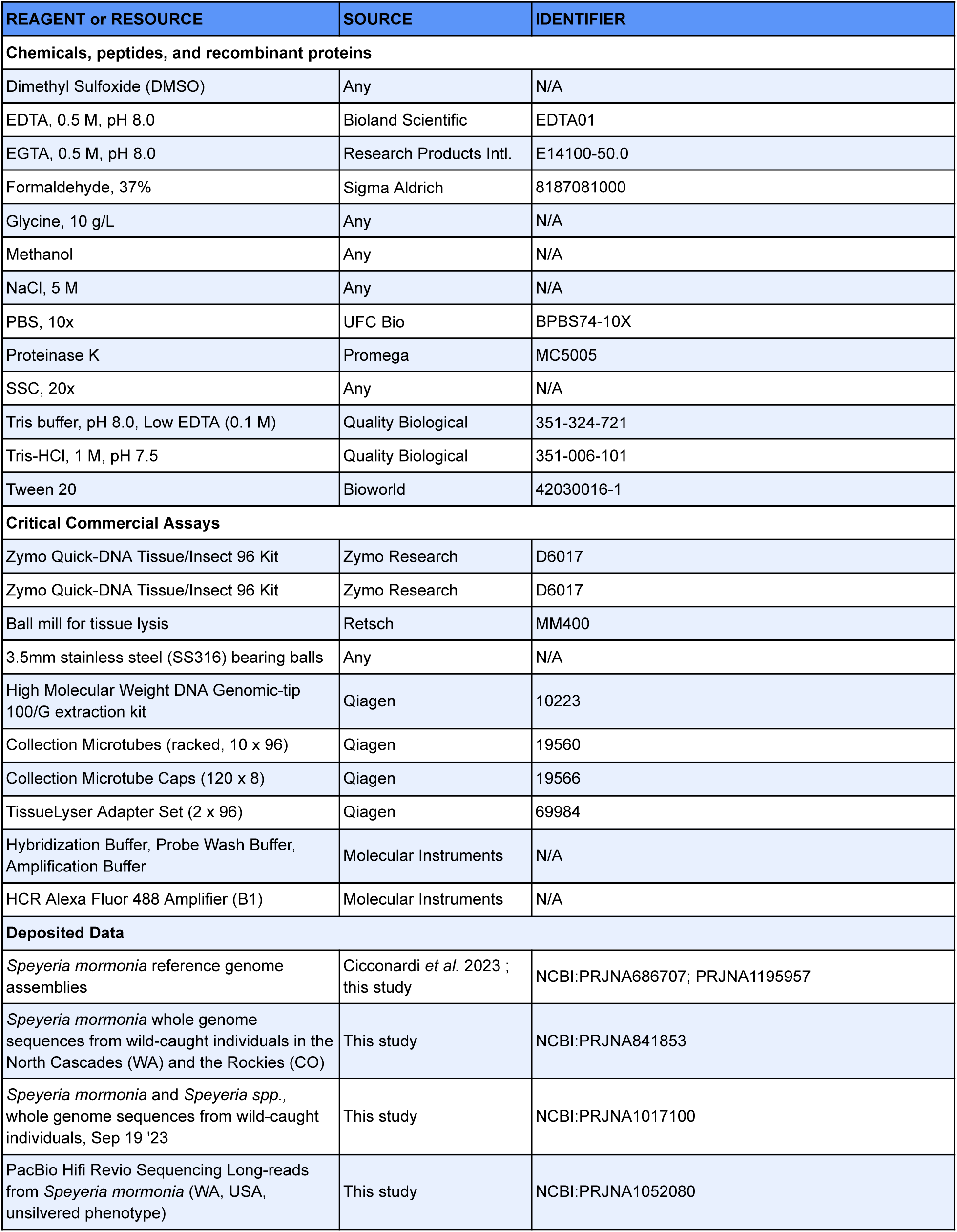
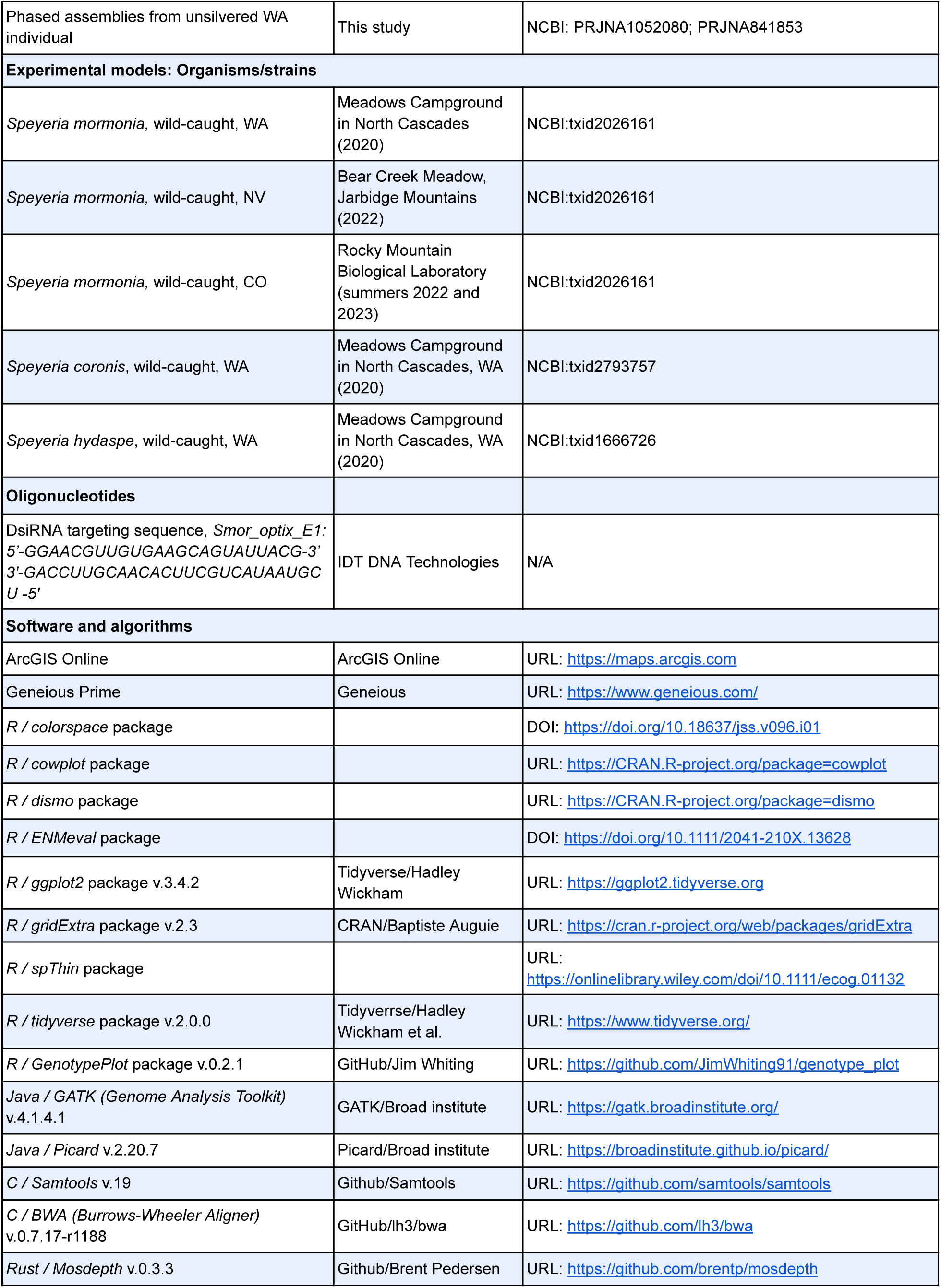
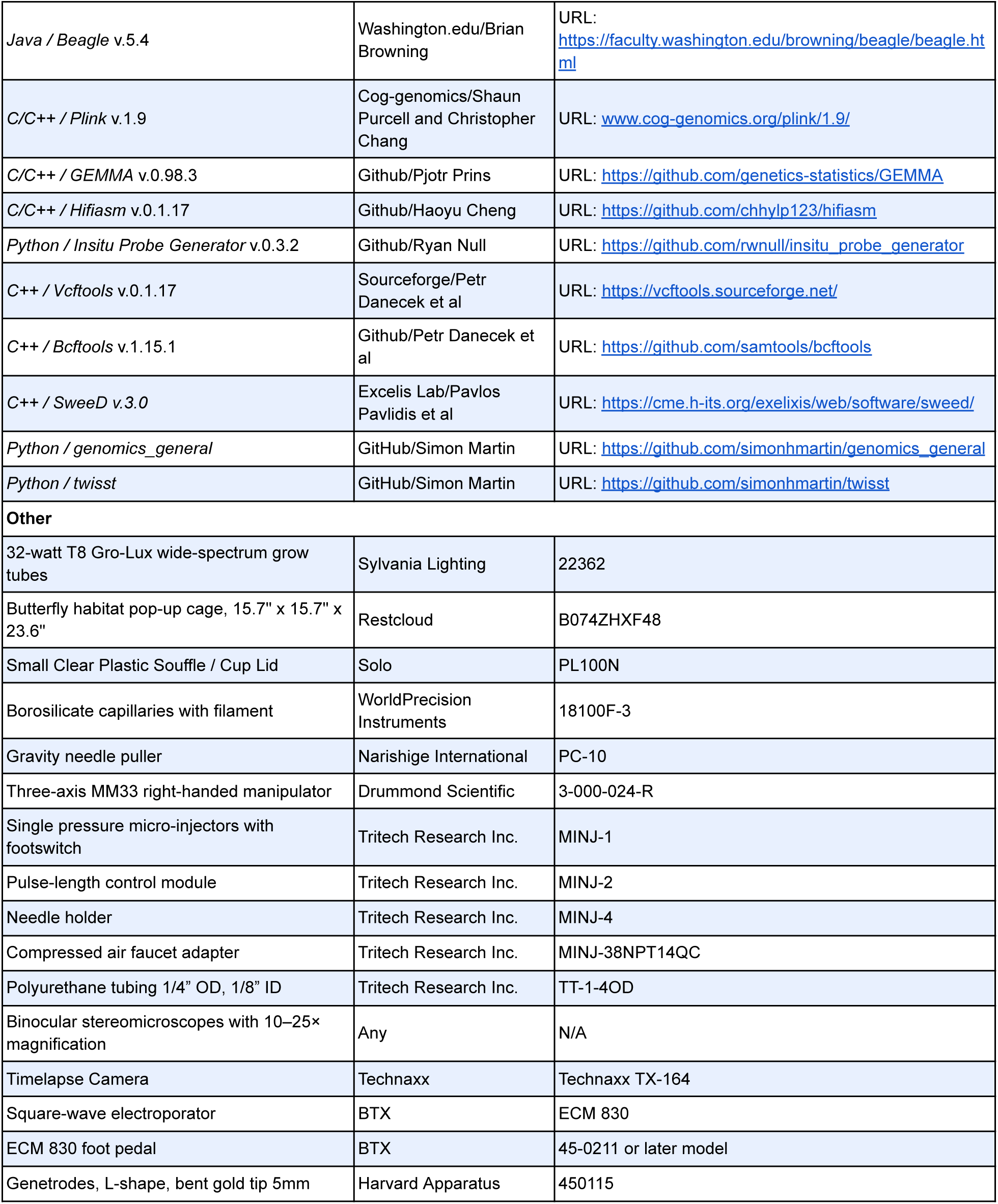
Key resources table.

### Resource availability

#### Lead contact

Further information and requests for resources should be directed to and will be fulfilled by the lead contact, Luca Livraghi. Any additional information required to reanalyze the data reported in this paper is available from the lead contact upon request.

#### Materials availability

This study did not generate new materials.

#### Data and code availability

The data and code used in this study were made available on the NCBI and GitHub repositories as detailed in the **Key Resources Table**.

### Experimental model and subject details

*Speyeria mormonia* (synonym: *Argynnis mormonia*) are the smallest of the North American fritillaries, occupying alpine meadows throughout the Western US and Canada. They are a univoltine species, diapausing as first instar larvae close to egg laying sites comprising several *Viola* species. Two populations were sampled for genome resequencing in this study, both polymorphic for the silvering trait. The first population was sampled in August 2020 from the Meadows Campground in the North Cascades, Washington. The outgroup species used in the study (*S. hydaspe* and *S. coronis*), were sampled from this same population. A second population of *S. mormonia* was sampled in August 2022 from Bear Creek Meadow in the Jarbidge Mountains, Nevada. Of note, the *Speyeria* genus was proposed to be revised as a sub-genus of the *Argynnis* genus, and the *Speyeria/Argynnis mormonia* species is undergoing a potential taxonomic revision, with a proposed reclassification of most of its subspecies in the species nova *Argynnis bischoffi*^48^. Given the widespread evidence of hybridization in this lineage, we are awaiting further phylogenomic studies to clarify its taxonomy.

## Method details

### Phenotypic data collection from butterfly collections and image repositories

A total of 9,760 occurrence records for *S. mormonia* were obtained from a combination of museum records, private collections, and online repositories (GBIF, iNaturalist), as summarized in **Table S1**. In brief, *S. mormonia* individuals were examined from museum and private collections, scoring the presence of the silvering trait for each locality. To supplement the silvering trait distribution, research-grade GBIF records including from iNaturalist entries featuring scorable pictures of ventral wing surfaces were added to the dataset. Unusual cases where ambiguous silvering traits existed were left unscored. Each individual and corresponding latitude and longitude, as well as silvering states where available, were plotted on a basemap in ArcGIS.

### Species distribution modeling

We used the occurrence records featured in **Table S1** to construct a suite of species distribution models (SDMs) for the species using WorldClim data and elevation as predictors^49^. Prior to modeling, we removed dubious records and those with erroneous coordinate data (*e.g.* occurring over bodies of water). We further thinned records by using a spatial buffer of 5 kilometers through the R package *spThin*^50^. Background points were randomly sampled across the extent of the study region. Using the package *ENMeval* we constructed MaxNet species distribution models using all combinations of linear, quadratic, and product predictors across regularization values of 0.5, 1, 2, and 3^51^. We partitioned the data using a checkerboard approach.

We assessed SDM performance by calculating the Akaike Information Criterion (AIC) score and selected the top performant model based on this metric. We also compared area under curve (AUC) and continuous Boyce index (CBI) scores, ensuring that these values were sufficiently robust in our top model. The top model achieved an AUC score of 0.92 and a CBI of 0.99 on the validation partition. We created a binary range map from this top model by thresholding such that occurrence probability values greater than 20% were represented as a potential range map. Finally, we clipped this binary map to an expert-drawn range map for the species^38^.

### Geospatial and bioclimatic regression analyses

Using the geographic phenotype data, we used beta-regression analysis to detect associations between local silvering frequency and environmental variables. To calculate silvering frequencies, phenotypic records were binned into 20-km radius clusters by calculating the distance between occurrence records from latitude and longitude coordinates, and applying a complete hierarchical clustering approach to these pairwise distances. Clusters of 20 km were identified by “cutting” the resulting dendrogram by this distance. Elevation data were obtained from the SRTM 90m Digital Elevation Database^52^. BioClim variables were obtained from WorldClim^49^. We obtained three additional variables – downward surface solar radiation, total precipitation, and leaf area index – from the Copernicus Climate Store ERA5-Land Monthly Dataset across the years 1970-2024^53^, for the adult flight period of the butterfly (June – September).

### Thermal emissivity properties of silver and unsilvered scales

To evaluate the impact of the silvering polymorphism on the thermodynamics of *Speyeria mormonia*‘s wings, we used thermal imaging to measure increase in wing temperature under sunlight illumination. **Figure S4A** illustrates the measurement setup modeling the thermodynamic system of a butterfly wing, as previously described^16^. Briefly, wings were suspended above the temperature-controlled cold plate which represents the cold sky, using a thermo-radiatively transparent polyethylene thin film. A xenon lamp (Thorlabs HPLS-30-03, color temperature = 6,500K) simulating the solar radiation illuminates the wing at an angle of 15° to the vertical. Although in the real world, the sun and sky are on the same side of the wing, which is different from our artificial system where the sun and sky are on opposite sides of the wing, the two thermodynamic systems are equivalent considering that the wing is extremely thin. Because of the thinness, the conduction through the wing thickness occurs instantly, resulting in an insignificant temperature gradient across the wing thickness. A thermal camera (FLIR T640) was positioned above the wing to monitor the thermal radiance from the wing along the normal axis. We first used the thermal camera to measure the optical properties of the wings. Since a butterfly wing is thin enough to be semi-transparent at the responsive wavelengths of the thermal camera, the irradiance perceived by the camera, Φ*_cam_*, comprises three components as shown in **Figure S4A**: (1) the thermal radiance from the wing, Φ*_w_*, (2) the thermal radiance of the ambient environment, Φ*_t_*, reflected by the wing and (3) the thermal radiance of cold plate underneath, transmitting through the wing, Φ*_r_*^16^. The perceived irradiance can be expressed as:

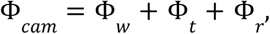

where:

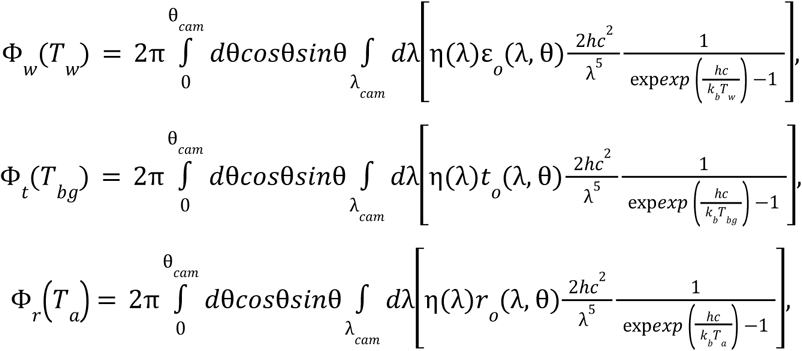

and where ε*_w_*, *t_w_*, *r_w_* are the emissivity, transmittance, and reflectance of the wing respectively. *T_w_*, *T_a_*, *and T_bg_* are the temperature of the wing, ambient environment, and cold plate. The resulting perceived irradiance is:

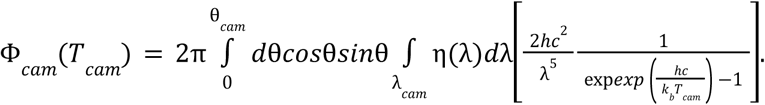

By applying a temperature difference between the ambient environment and the cold plate, the temperature of the wing measured by the camera, *T*_*cam*_, would range from *T*_*bg*_ to *T*_*a*_, dependent on thickness. To simplify these relations, we made a few approximations. First, we assumed the wings’ reflectance to be small and nearly constant, using a reflectance value of 0.05 in calculations. The transmission was then calculated as:

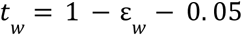

Secondly, when the xenon lamp was off, the true wing temperature, *T*_*w*_, was approximated as the ambient temperature of the ambient temperature, given the wing was sufficiently distant from the cold plate. These assumptions simplified the equations to:

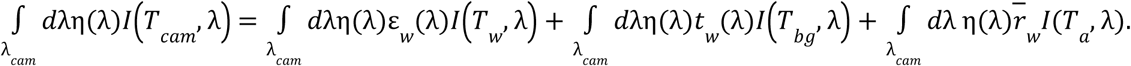

By capturing a single snapshot of the wing with the xenon lamp off, we can derive the optical properties of interest using the known temperatures of the ambience and cold plate. After acquiring the approximated emissivities of the wings, we turned on the lamp and recorded the rise of wing temperature. The measured temperatures were then converted into the real wing temperatures following the same equation with the previously measured optical properties.

The illumination of the Xenon lamp had an intensity of 60mW/cm^2^, which is about 2/3 full sun. We measured three pairs of unsilvered/silver morphs of *Speyeria mormonia* to compare the temperature rises of the patches at the same locations of their hindwings (**Figure S4B**) induced by the illumination.

### Speyeria mormonia crosses

Thirty-one silver and 23 unsilvered wild-caught females were transported in individual glassine envelopes to the lab for egg laying. Females were housed in plastic containers lined with filter paper. Infrared heat lamps supplied high temperature and light, which stimulate egg-laying^54^. For the original wild-caught females, lamps were operated by an automatic timer on a 17L:7D photoperiod. For later generations, the light period was adjusted to 14L:10D. Temperature ranged from 22 to 30°C and relative humidity ranged from 74 to 89%. All adults were fed with a 20% sucrose solution in an artificial feeder. Females were provided with either *Viola sororia* or *V. papilionacea* leaves for egg laying, which were briefly soaked in a mild bleach solution and placed in a water-pik. Upon hatching, first instar larvae were transferred at 2-7°C for the diapause period. Larval diapause was interrupted by transferring the larvae to a moist filter paper, and subsequently transferring single broods to individual rearing containers with *V. papilonacea* leaves. Larvae were allowed to pupate, given a unique code, and identified to a brood and larval cohort. Silvering trait was recorded and used to establish crosses over two subsequent generations. Phenotypic proportions and inferred genotypes for broods with more than 3 individuals are reported in **Table S2.**

### Wild population sampling and DNA extraction

Individuals sampled for genome re-sequencing included 19 unsilvered and 20 silver individuals from the Meadows Campground in the North Cascades, WA and 15 unsilvered and 16 silver individuals from the Bear Creek Meadow in the Jarbidge Mountains, Nevada (Table Sx). Wild-caught samples were collected, wings stored in glassine envelopes and bodies kept in DESS buffer (20% dimethyl sulfoxide, 0.25M ethylenediaminetetraacetic acid, saturated sodium chloride, pH 8.0). Samples were subsequently stored at –20°C. DNA was extracted from adult thoraxes using a Zymo Quick-DNA Tissue/Insect 96 Kit and concentrations measured with a Qubit spectrophotometer.

### Genome sequencing and genotype calling

Sequencing libraries (PE 150) were generated and sequenced in an Illumina NovaSeq6000 S4 Flow cell at the Institute for Genome Sciences (University of Maryland – Baltimore) generating resequencing data at 19.7x mean coverage, as measured with the *mosdepth* package^55^. Each sample was then aligned to the *S. mormonia* reference assembly^8^ using *bwa-mem* with default parameters^56^. PCR duplicates were marked and removed using *picard*^57^. Genotypes were called using the Genome Analysis Tool Kit (GATK) *HaplotypeCaller*^58^ using parameters optimized for butterfly systems^5^. Briefly, joint genotyping of individual genomic VCF records (gVCF) was performed with GATK’s *GenotypeGVCFs*, using default parameters. Genotype calls were included in downstream analysis only if they met the following criteria: quality (QUAL) ≥ 30, minimum depth ≥ 10, maximum depth ≤ 100 (to avoid false SNPs due to mapping in repetitive regions), overall depth ≤ 100 × number of samples, strand bias (FS) < 200, quality by depth (QD) ≥ 5, and for variant calls, genotype quality (GQ) ≥ 30. The resulting VCF files were split into indel and SNP files, and subsequent analysis were performed only on SNPs. Further filtering was performed using *bcftools*^59^, removing minor allele frequencies (MAF) of > 0.02 and missing genotypes (F_MISSING) of < 0.1.

### Genome-Wide Association Study, Principal Component Analysis and SNP visualization

To perform the GWAS, VCF files were further filtered by removing non-variant sites and keeping only bi-allelic SNPs. Missing genotypes were imputed and phased using *beagle* v5.4^60,61^. A binary fileset was generated using *PLINK* v1.9^62^ for input into *GEMMA* v0.98.3^63^, which was used to compute site-wise Wald χ^2^ test *P*-values, accounting for population structure by supplying a relatedness matrix as a covariate. Genome-wide associations were performed both separately for each population (WA and NV) as well as combining both datasets. To perform a principal component analysis on the whole genome dataset, SNPs were first linkage-pruned from a VCF file containing samples from both populations, then a PCA was performed on both a VCF file containing SNPs along the whole genome, as well as local sets of SNPs confined to the Genome Wide Association interval. PCA analyses were performed using *PLINK* v1.9^62^. The package *genotypeplot*^64^ was used to inspect the GWA interval for trait associated SNPs, plotting each SNP coded in the VCF file as either homozygous for the reference allele, homozygous for the alternate allele, or heterozygous.

### Long-read genome assembly of an unsilvered morph

High-molecular-weight DNA was extracted from an unsilvered individual from the WA population, homozygous for the fixed associated SNPs within the GWA interval (**Figure 2D**), using the QIAGEN Genomic-tip 100/G kit. DNA was sequenced on a PacBio Revio system at the Institute for Genome Sciences (University of Maryland – Baltimore), producing 28.2 Gb of sequence data. Genome assembly was performed using *HiFiasm* v0.15.1^65^ using default parameters, resulting in two phased haploid assemblies with lengths of 447.9 Mb (N50 = 15.4 Mb; 201 scaffolds) and 442.8 Mb (N50 = 14.4 Mb; 121 scaffolds).

### Rearing of *Speyeria mormonia* for fluorescent *in situ* hybridization and RNA interference

First-instar larvae from a population at the Rocky Mountain Biological Laboratory in Colorado (>99.9% silver^20^), were brought out of diapause by transferring them from moist glass scintillation vials to tupperware containing leaves of *V. sororia* and exposing them to constant light for 72hrs at 25°C. Larvae were regularly stimulated by gentle stimulation using a fine paintbrush, or by lightly blowing on them while monitoring for signs of feeding behavior, upon which they were transferred to a 18L:6D photoperiod and kept at 25°C. Larval rearing containers consisted of vented tupperwares^66^ with a folded Kimwipe at the bottom, with *V. sororia* leaves tightly inserted in 1.5-mL tubes filled with shipping refrigerant gel to limit water evaporation. Total pupation time was recorded from onset of pupariation formation to eclosion using a TX-164 timelapse camera (Technaxx). Total development time was used to calculate pupal % development for accurate timings of dissections.

### Hybridization chain reaction (HCR) in *S. mormonia* pupal wings

Ten HCR^67^ probes were designed against the *optix* coding exon (**Table S3**), using the *insitu_probe_generator* tool^68^, permitting a GC content in the 35-75% range, and allowing limiting runs of poly-GC and poly-AT to a maximum of 3 bp. Pupal wings were dissected from the pupal case in cold 1X PBS as previously described in other butterflies^3^, transferred to a fixative solution (750 μL PBS 2 mM EGTA, 250 μL 37% formaldehyde) containing 9.25% formaldehyde at room temperature for 30 min and washed four times in PBS containing 0.01% Tween20 (PBT). For pupae at earlier stages of development (28%), samples were permeabilized in 1 μg/μL of ProteinaseK diluted in PBT solution for 5 min. Later-stage pupae (54%) were first permeabilized using detergent solution (10% SDS, 20% Tween, 1 M Tris-HCl, pH 7.5, 0.5 M EDTA, pH 8.0, 5 M NaCl) for 30 min at room temperature and then further digested in 1 μg/μL of Proteinase K in PBT for 7 min at 55°C. Samples were then washed with a stop solution containing PBT and 2-mg/mL glycine and followed by two additional PBT washes. After transferring wings to a post-fix solution (850 μL PBT, 150 μL 37% formaldehyde) containing 5.55% formaldehyde for 20 min, wings were washed four times with PBT before following the rest of the protocol as in previously published procedures^69^.

### Pupal wing electroporation and RNA interference

Dicer-substrate siRNAs (DsiRNAs) were designed against the *optix* coding exon (5’-GGAACGUUGUGAAGCAGUAUUACG-3’), ordered as Custom DsiRNAs with standard purification (IDT DNA Technologies), resuspended at 100 μM in 1X *Bombyx* injection buffer (pH 7.2, 0.5 mM NaH_2_PO_4_, 0.5 mM Na_2_HPO_4_, 5 mM KCl), and stored as frozen aliquots at –70°C until use. Electroporation procedures followed a previously established protocol^21^. To target the dorsal pupal wings, the negative electrode was placed in contact with a droplet of phosphate buffered saline (PBS) positioned on the ventral side of the peeled wing, while the positive electrode was placed on the agarose pad on the dorsal side (poles are reversed when targeting the ventral wing surface). Electroporation was then carried out with 5 square pulses of 280 ms at 10 V, separated by 100-ms intervals.

### Calculation of silver locus allele frequency for population genetic summary statistics and signatures of selection

To calculate haplotype-specific statistics, we first phased a VCF file containing all populations using *beagle* v5.4^60,61^. A custom script (https://github.com/MilesLuca/Speyeria_Silver) was then used to separate each haplotype into a new entry in the VCF, recoding individual haplotypes as either “_A” or “_B” based on the three diagnostic SNPs identified in the GWAS. Fixation index (Fst), absolute divergence (Dxy), nucleotide diversity (π), Tajima’s D and proportion of introgression (*f_d_*) statistics were then calculated on individual alleles using the genomics_general toolkit (https://github.com/simonhmartin/genomics_general). The same phased and recoded VCF was used to generate selective sweep statistics using the software package *SweeD*^22^. Support for phylogenetic hypotheses consistent with shared haplotypes of unsilvered alleles were evaluated using Twisst^25^ (topology weighting by iterative sampling of subtrees; https://github.com/simonhmartin/twisst).

## Author contributions

Conceptualization, LL, JJH, WMR, CLB and AM; Methodology, LL, VMS, CCT, NY; Investigation, LL, JJH, LSL, VMS, CCT, NY, SMVB, AH, CMTK, WMR, CLB, and AM; Writing, LL, JJH and AM; Funding Acquisition and Resources, LL and AM.

## Supplemental information

**Figure S1.**
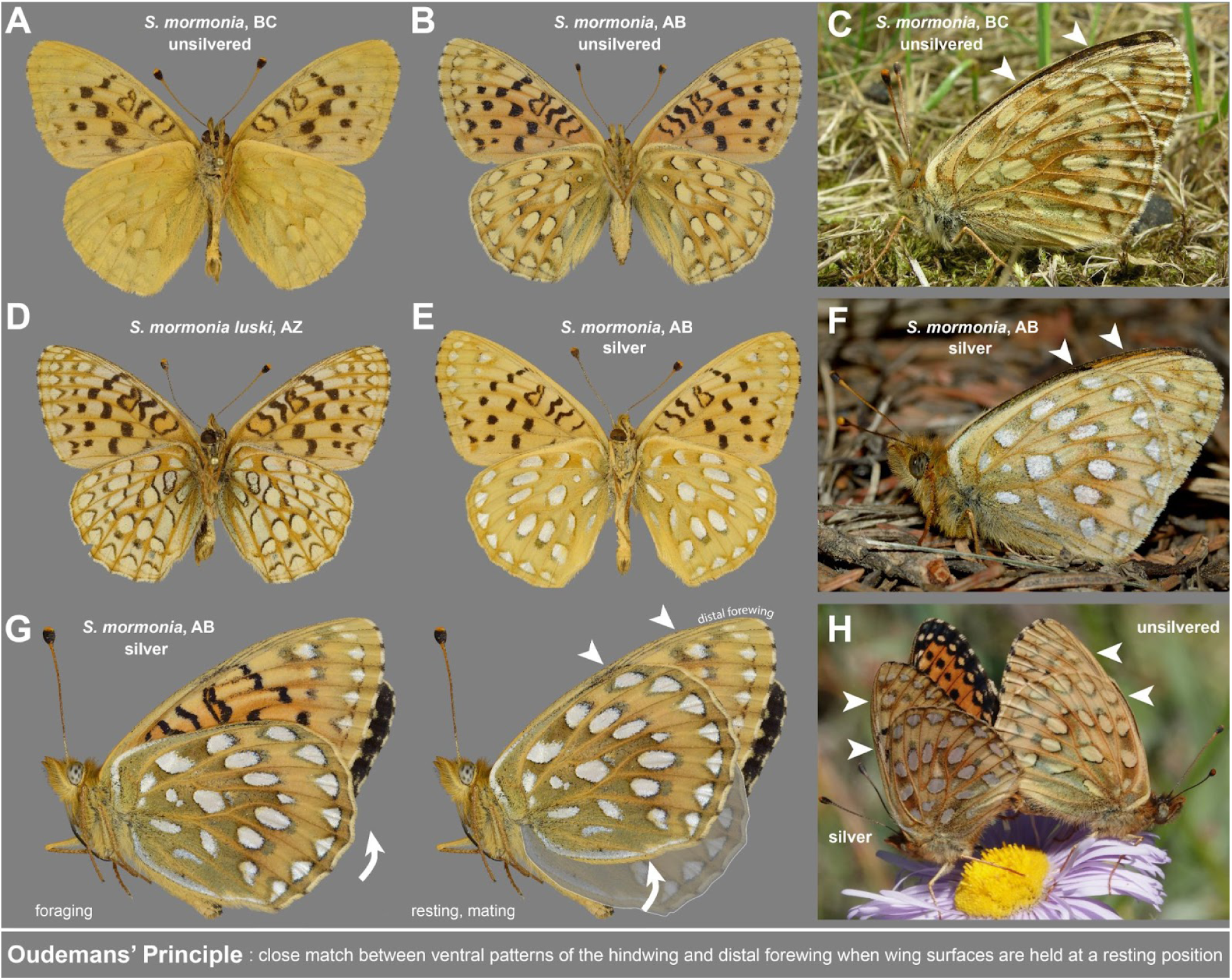
Silver dimorphic patterns are exposed in the resting position. *S. mormonia* butterflies show a wide range of variation in their ventral wing surfaces including unsilvered (**A-D**) and silver morphs (**E-F**). In the forewing, these variations are restricted to spots of the distal forewing. At rest and during mating (**G-H**), the spotted areas of the two align and match (arrowheads), while the more conspicuous black and orange patterns of the forewing are covered by the hindwing. This phenomenon follows the Oudemans principle^43^ and suggests a function in camouflage at rest. All photographs specimens were contributed by Norbert G. Kondla from his collection. More extensive documentation of phenotypic variations within the genus are available on his online photo album (URL: https://flic.kr/s/aHskjkQSPr). Photographs were not reproduced at scale and include the following specimen locations. **(A)** Collected Good Hope Lake (BC), 1995-07-11. **(B)** Collected near Gloomy Creek Rd (AB), 2015-08-12. **(C)** Photographed near Rossland (BC), 2007-07-23. **(D)** Collected at Paradise Creek (White Mountains, AZ), 2001-07-28. **(E)** Collected at Whiskey Gap (AB), 1983-06-23. **(F)** Photographed near Limestone Mountain (AB), 2016-08-17. **(G)** Collected near Moose Mountain (AB), 2023-06-24. The hindwing was digitally rotated in the right panel to recreate the resting position. **(H)** Heteromorphic mating pair photographed at the Porcupine Hills (AB), 2022-08-07.

**Figure S2.**
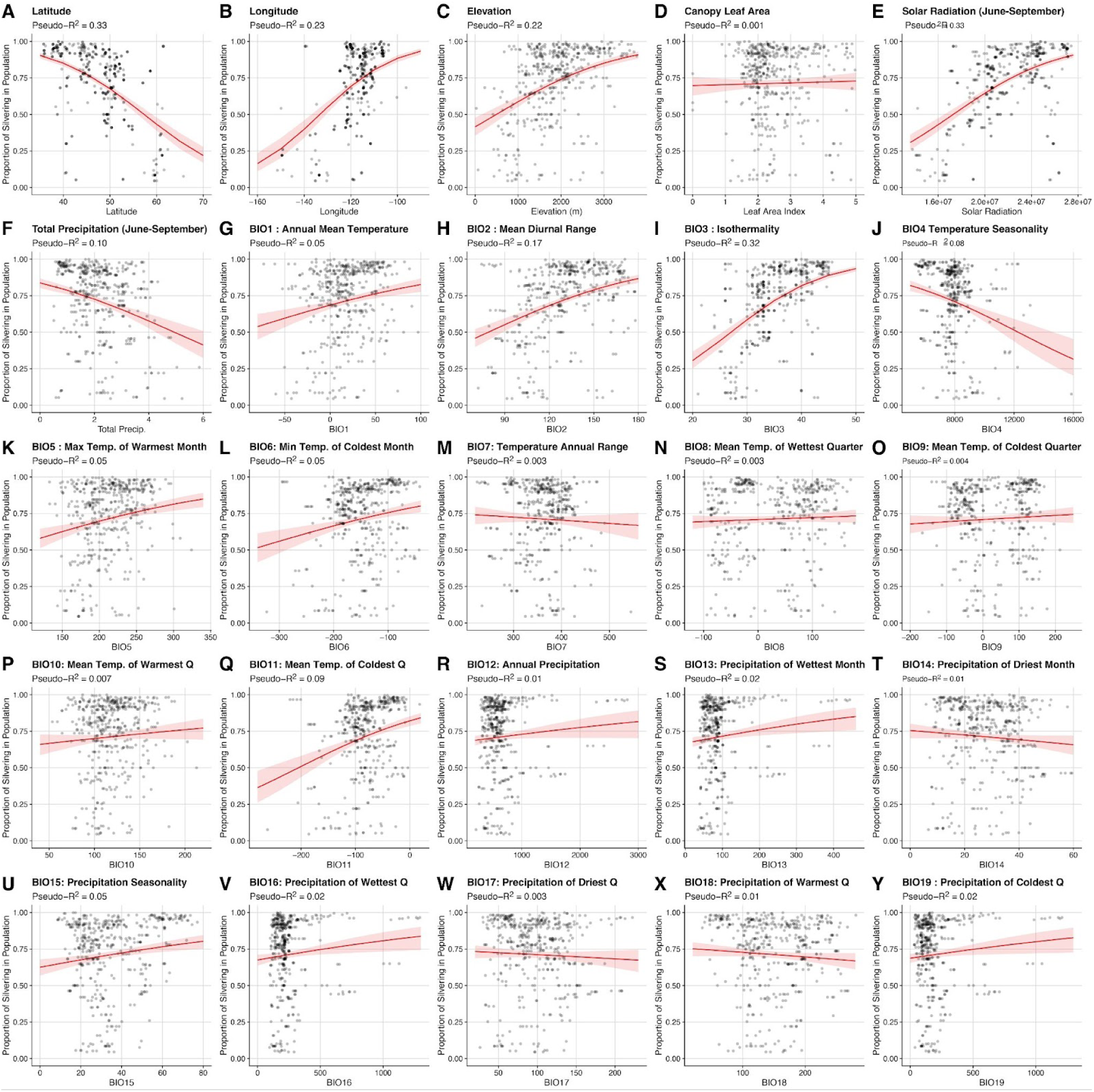
Beta-regression analysis of silvering proportions against geographic and bioclimatic variables. Dots feature silvering proportions (Table S1) following aggregations of N>5 individuals with a radius of 20 km, and the corresponding independent variables at these locations. Regression lines are shown in red. **A-C**. Latitude, longitude and imputed elevation. **D.** Canopy leaf area. **E-F.** Solar radiation and Cumulative precipitation over the June-September months (maximum adult flight period for *S. mormonia*). **G-Y.** BioClim variables.

**Figure S3.**
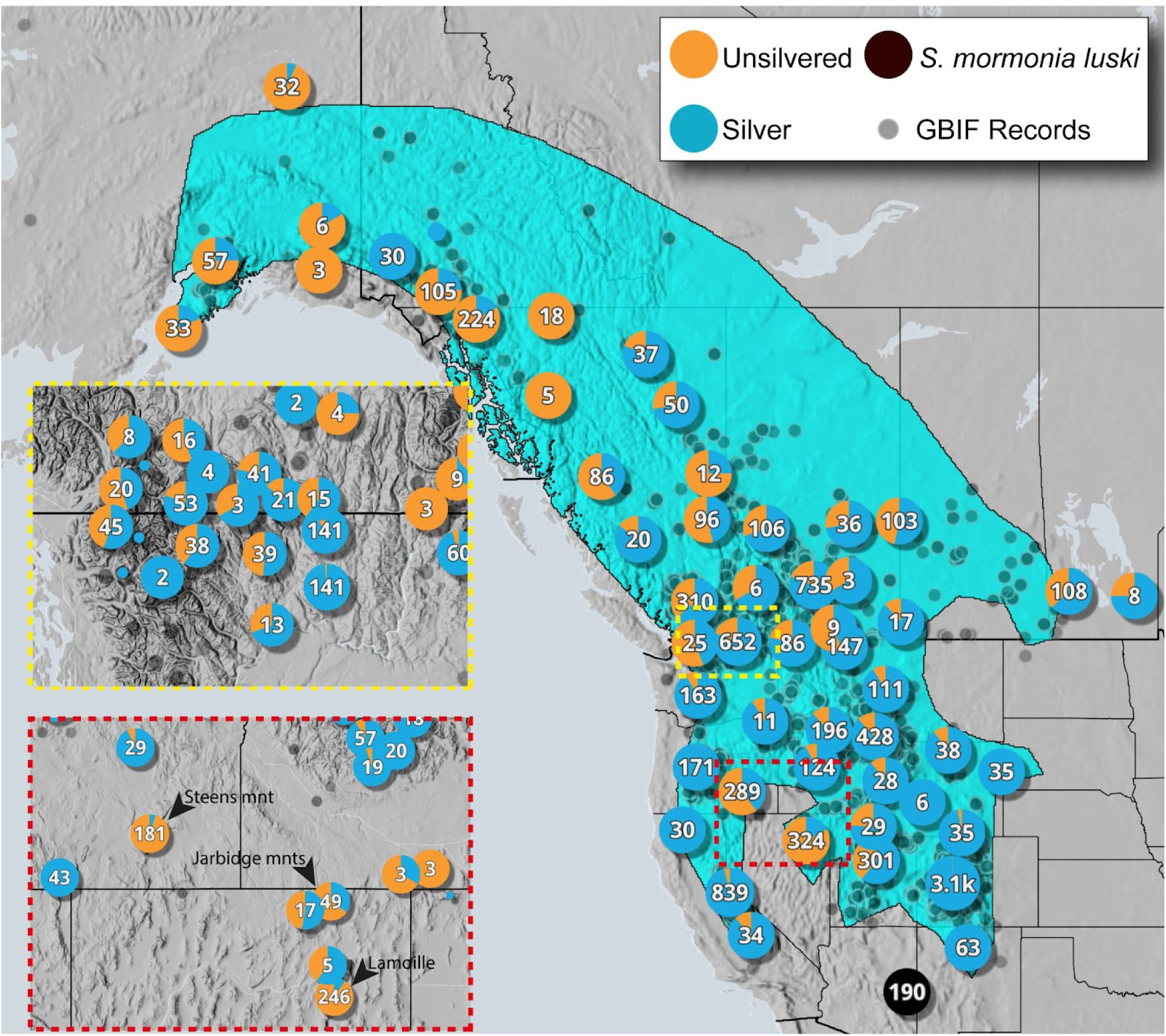
Zoomed-in views of silvering polymorphism in sampled regions of Washington (WA) and Nevada (NV). The distribution of silver (blue) and unsilvered (brown) phenotypes in *S. mormonia* populations is shown with a closer focus on the regions sampled in this study. The upper inset (yellow border) highlights the North Cascades region in Washington (WA), while the lower inset (red border) focuses on the predominantly unsilvered population in the Steens Mountains (South East Oregon), and the Jarbidge and Ruby Mountains in Northern Nevada (NV). Phenotype frequencies are represented by pie charts, with the number of individuals displayed in each population. Gray circles represent additional *S. mormonia* occurrence records from the Global Biodiversity Information Facility (GBIF) where phenotypic data was not available. A distinct subspecies, *S. mormonia luski*, found in Arizona and exhibiting an atypical phenotype (**Figure S1D**), is shown in black. A species distribution model imputed from bioclimatic modeling is outlined in cyan.

**Figure S4.**
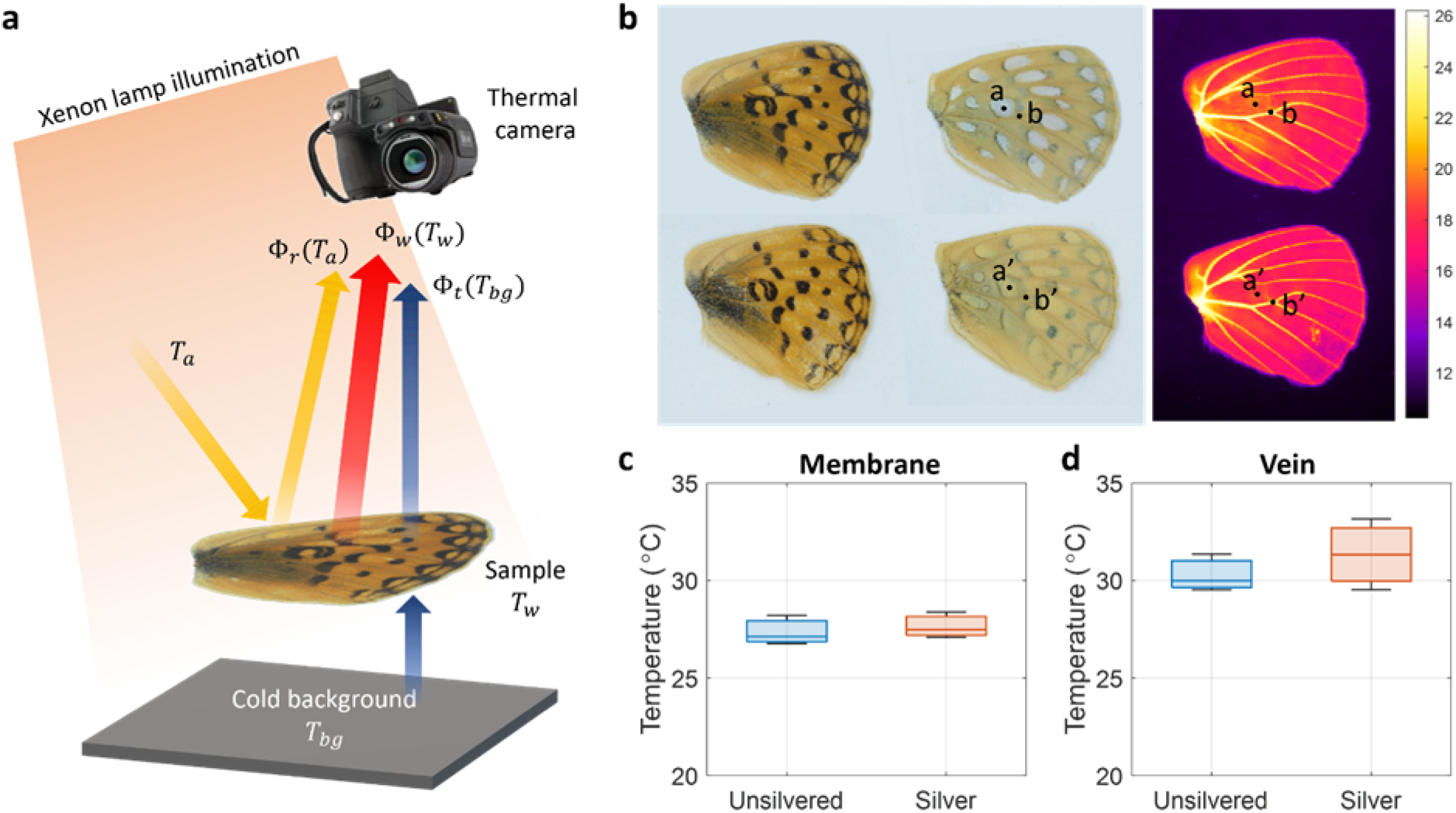
No difference in thermal emissivity between silver and unsilvered spots. **(A)** Schematics of the experimental setup to measure the equilibrium temperature under xenon illumination levels approximating 2/3 of full sun conditions. The configurations simulate the thermodynamic system of a butterfly wing in nature. **(B)** A pair of hindwings of the silver (upper row) and unsilvered (lower row) morphs of *Speyeria mormonia*. Left and middle columns: optical images of the wing dorsal and ventral sides. Right column: thermal image of the wings taken by the setup in **(A)** with the illumination of xenon lamp on. Temperatures were measured at the center of a silver/unsilvered patch on the end of the discal area (a, a’), and on the adjacent discal crossvein (b, b’). **(C)** Equilibrium temperatures of silver/unsilvered patches (a and a’; N = 3). **(D)** Equilibrium temperatures of the adjacent discal crossveins (b and b’; N = 3). Both differences are non significant; membrane (c) *t*-test *p*-value = 0.6514; vein (d) *t*-test *p*-value = 0.4417.

**Figure S5.**
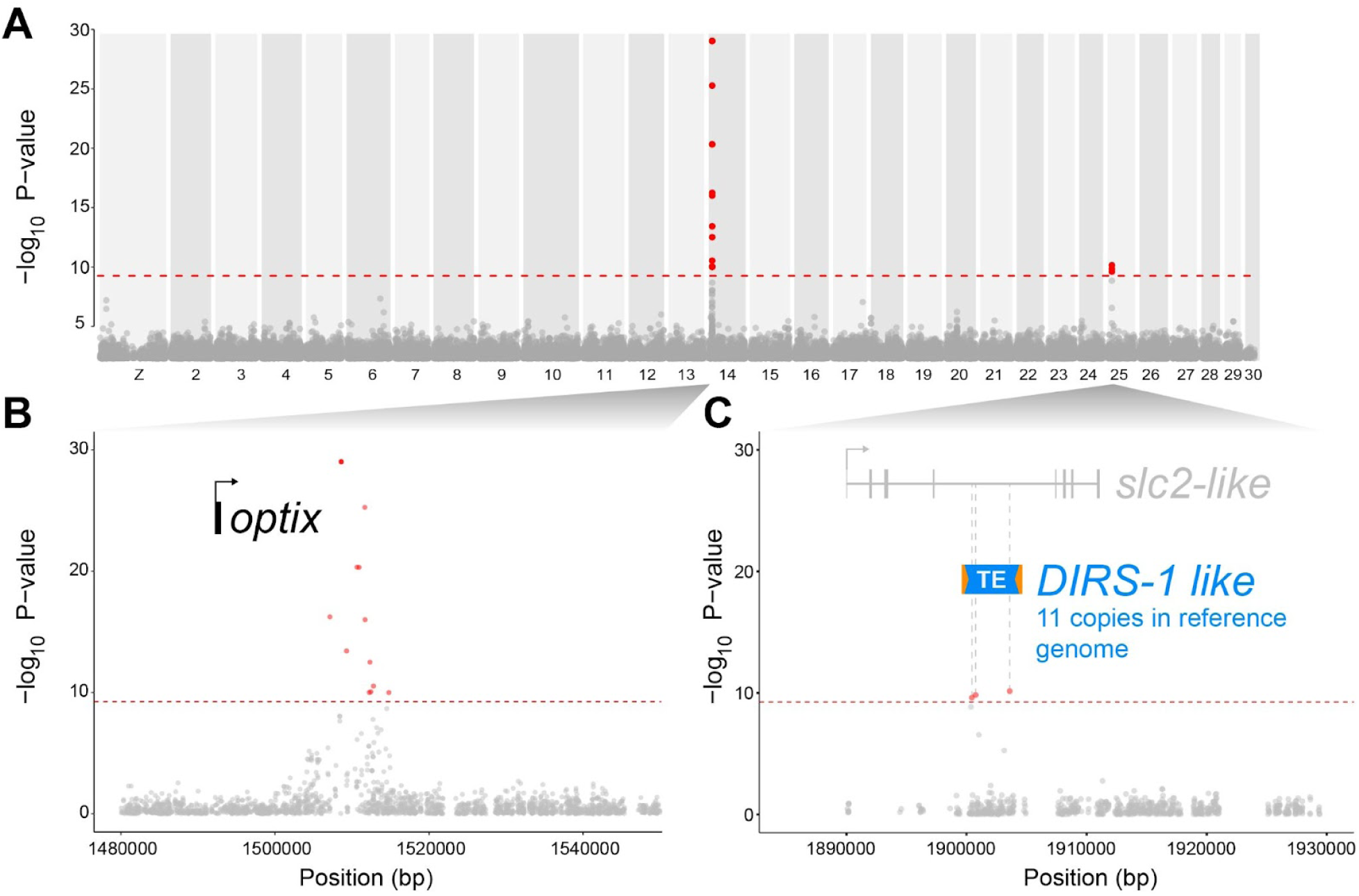
Genome-wide association study (GWAS) results for silvering for a combined dataset of both WA and NV populations. **(A)** Manhattan plot for GWAS analysis of combined datasets from WA and NV. The red line indicates the Bonferroni-corrected significance threshold (P < 0.01). Significant loci surpassing this threshold are highlighted in red. **(B)** GWAS SNPs for silvering map to the 3’ region of the *optix* transcription factor gene. **(C)** A secondary GWA signal corresponds to 3 SNPs detected exclusively in the NV population (**Figure 2A**), that map inside an inactive transposable element of the *DIRS1-like* family. We found 11 copies of this 5-kb element in the reference genome, and SNPs in the center of this region fail to form a continuous associated haplotype, raising the possibility that the GWA signal is spurious or unspecific to chromosome 25. Blue: *DIRS-1* transposon region. Orange: inverted terminal repeats.

**Figure S6.**
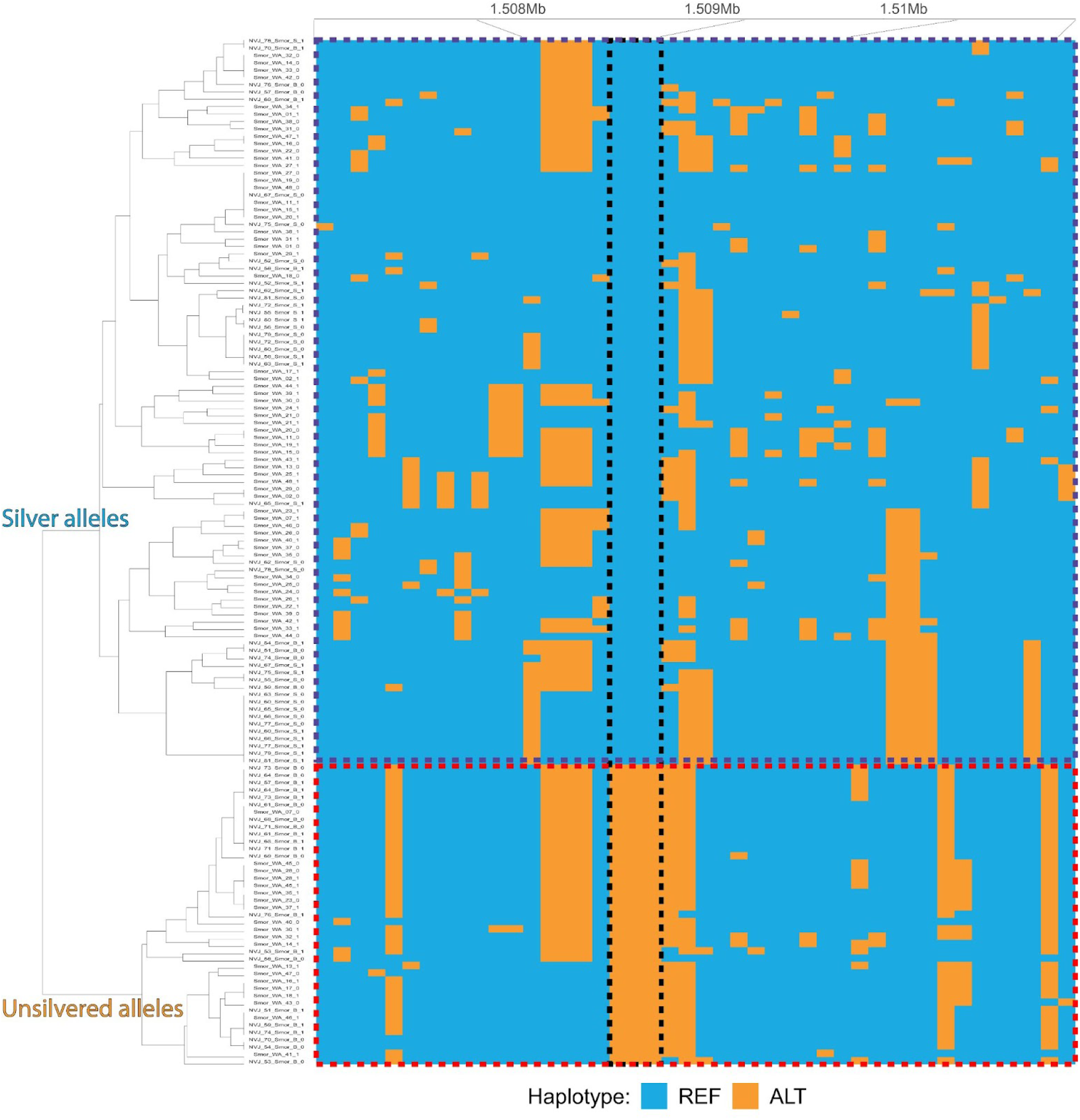
Genotype plot of the GWAS region clustered by phased haplotypes. Each row represents an individual, and each column shows a color-coded SNP, with alleles coded as either reference (homozygous for silver) or alternative. The cladogram on the left illustrates sample clustering using a PCA method implemented in *Genotype_Plot*, and is not indicative of allele phylogeny. Presumptive haplotypes associated with the unsilvered phenotype are highlighted with a red dashed box, while those associated with the silver phenotype are highlighted with a blue dashed box. The vertical black dashed box highlights the 3 SNPs that are fixed between silver and unsilvered haplotype groups.

**Figure S7.**
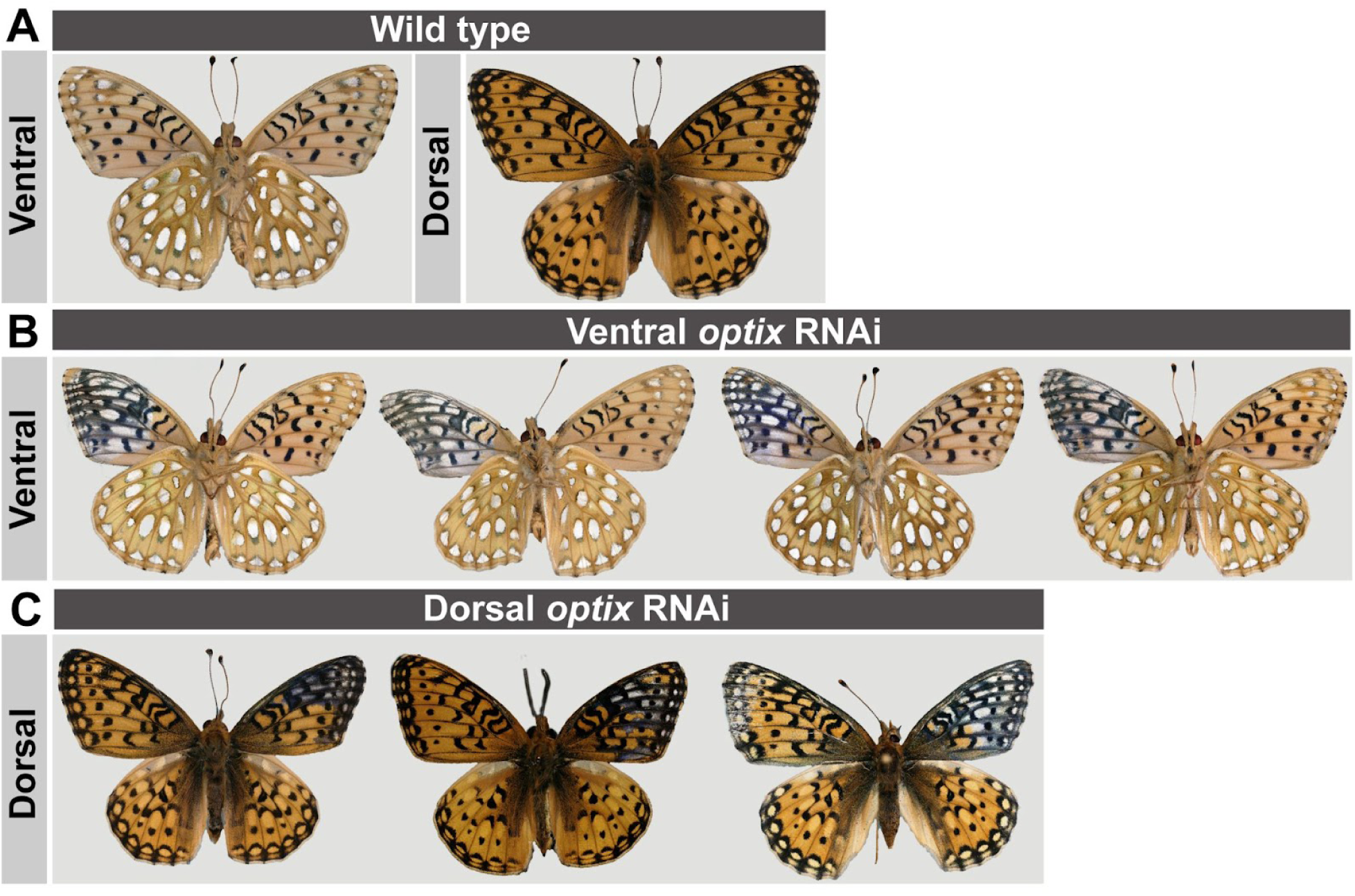
Electroporation mediated RNA interference (RNAi) against *optix* in pupal wings results in a transformation of unsilvered/orange scales to silver. **(A)** Wild-type dorsal and ventral *S. mormonia* wing patterns, contrasted with *optix* RNAi treated forewings on the ventral **(B)** and **(C)** dorsal wing surfaces. RNAi electroporation was systematically targeting the right wing, leaving the left sides as contralateral controls. Targeting of the ventral vs. dorsal sides is performed using electrode polarity, with the (+) electrode positioned on the side of the targeted surface^70^.

**Table S1.** Geographic distribution of silver dimorphism in *S. mormonia* in museum and online image repositories. This dataset is provided online as a supplementary Excel file. Related to **Figure 1E**.

**Table S2.** Frequencies of silvering phenotypes of captive-reared *S. mormonia* individuals. Each row gives the frequencies for a single brood from a phenotypic cross of the type listed in the first column, with probable parental genotypes for each cross, as inferred from parental phenotypes, phenotypes of grandparents, and offspring phenotype frequencies. Generation 1 refers to offspring of the original, wild-caught females. Broods that yielded < 4 individuals are omitted. For broods showing mixed phenotypes, results of ꭓ² tests against expected proportions based on the inferred parental genotypes are presented. Fisher exact tests indicate a lack of association between silvering frequency and sex for mixed broods. Abbreviations: S = silver phenotype; U = unsilvered phenotype; *U* = unsilvered-dominant allele; *s* = silver-recessive allele; – = either *s* or *U*.

| Parental phenotypes<br>(♀ × ♂) | Inferred parental genotypes | Generation | Females |  | Males |  | Sex linkage<br>Fisher exact *p* | Both sexes |  |  |  |
| --- | --- | --- | --- | --- | --- | --- | --- | --- | --- | --- | --- |
| | | | % S | N | % S | N | | % S | N | $\chi^2$ | *p* |
| S × unknown | ss × *U*- | 1 | 0 | 3 | 0 | 2 |  | 0 | 5 |  |  |
| S × unknown | ss × *Us* | 1 | 22.2 | 9 | 69.2 | 13 | **0.081** | 50 | 22 | 0 | **1** |
| S × S | ss × ss | 2 | 100 | 48 | 100 | 59 |  | 100 | 107 |  |  |
| S × S | ss × ss | 3 | 100 | 5 | 100 | 3 |  | 100 | 8 |  |  |
| S × S | ss × ss | 3 | 100 | 22 | 100 | 26 |  | 100 | 48 |  |  |
| S × S | ss × ss | 3 | 100 | 28 | 100 | 28 |  | 100 | 56 |  |  |
| S × S | ss × ss | 3 | 100 | 14 | 100 | 23 |  | 100 | 37 |  |  |
| S × S | ss × ss | 3 | 100 | 26 | 100 | 25 |  | 100 | 51 |  |  |
| S × S | ss × ss | 3 | 100 | 27 | 100 | 39 |  | 100 | 66 |  |  |
| S × S | ss × ss | 3 | 100 | 7 | 100 | 8 |  | 100 | 15 |  |  |
| S × U | ss × *Us* | 3 | 60 | 5 | 87.5 | 8 | **0.511** | 76.9 | 13 | 3.77 | **0.052** |
| S × U | ss × *Us* | 3 | 66.7 | 3 | 66.7 | 3 | **1** | 66.7 | 6 | 0.67 | **0.414** |
| S × U | ss × *Us* | 1 | 57.9 | 19 | 44.4 | 18 | **0.517** | 51.4 | 37 | 0.03 | **0.869** |
| S × U | ss × *Us* | 3 | 50 | 8 | 45.5 | 22 | **1** | 46.7 | 30 | 0.13 | **0.715** |
| U × S | *Us* × ss | 1 | 60 | 5 | 75 | 8 | **1** | 69.2 | 13 | 1.92 | **0.166** |
| U × S | *Us* × ss | 3 | 62.5 | 8 | 57.14 | 7 | **1** | 60 | 15 | 0.6 | **0.439** |
| U × S | *Us* × ss | 2 | 66.7 | 21 | 55.6 | 36 | **0.577** | 59.7 | 57 | 2.12 | **0.145** |
| U × S | *Us* × ss | 2 | 62.5 | 8 | 35.3 | 17 | **0.389** | 44 | 25 | 0.36 | **0.549** |
| U × S | *Us* × ss | 3 | 52.2 | 23 | 35.1 | 37 | **0.282** | 41.7 | 60 | 1.67 | **0.197** |
| U × S | *UU* × ss | 3 | 0 | 66 | 0 | 60 |  | 0 | 126 |  |  |
| U × U | *Us* × *Us* | 2 | 0 | 1 | 50 | 4 | **1** | 40 | 5 | 0.6 | **0.439** |
| U × U | *Us* × *Us* | 1 | 44.4 | 9 | 25 | 12 | **0.397** | 33.3 | 21 | 0.78 | **0.378** |
| U × U | *Us* × *Us* | 2 | 0 | 1 | 14.3 | 7 | **1** | 12.5 | 8 | 0.67 | **0.414** |
| U × U | *U*- × *UU* or reverse | 3 | 0 | 4 | 0 | 2 |  | 0 | 6 |  |  |
| U × U | *U*- × *UU* or reverse | 3 | 0 | 7 | 0 | 9 |  | 0 | 16 |  |  |
| U × U | *Us* × *UU* | 2 | 0 | 22 | 0 | 56 |  | 0 | 78 |  |  |

**Table S3.**
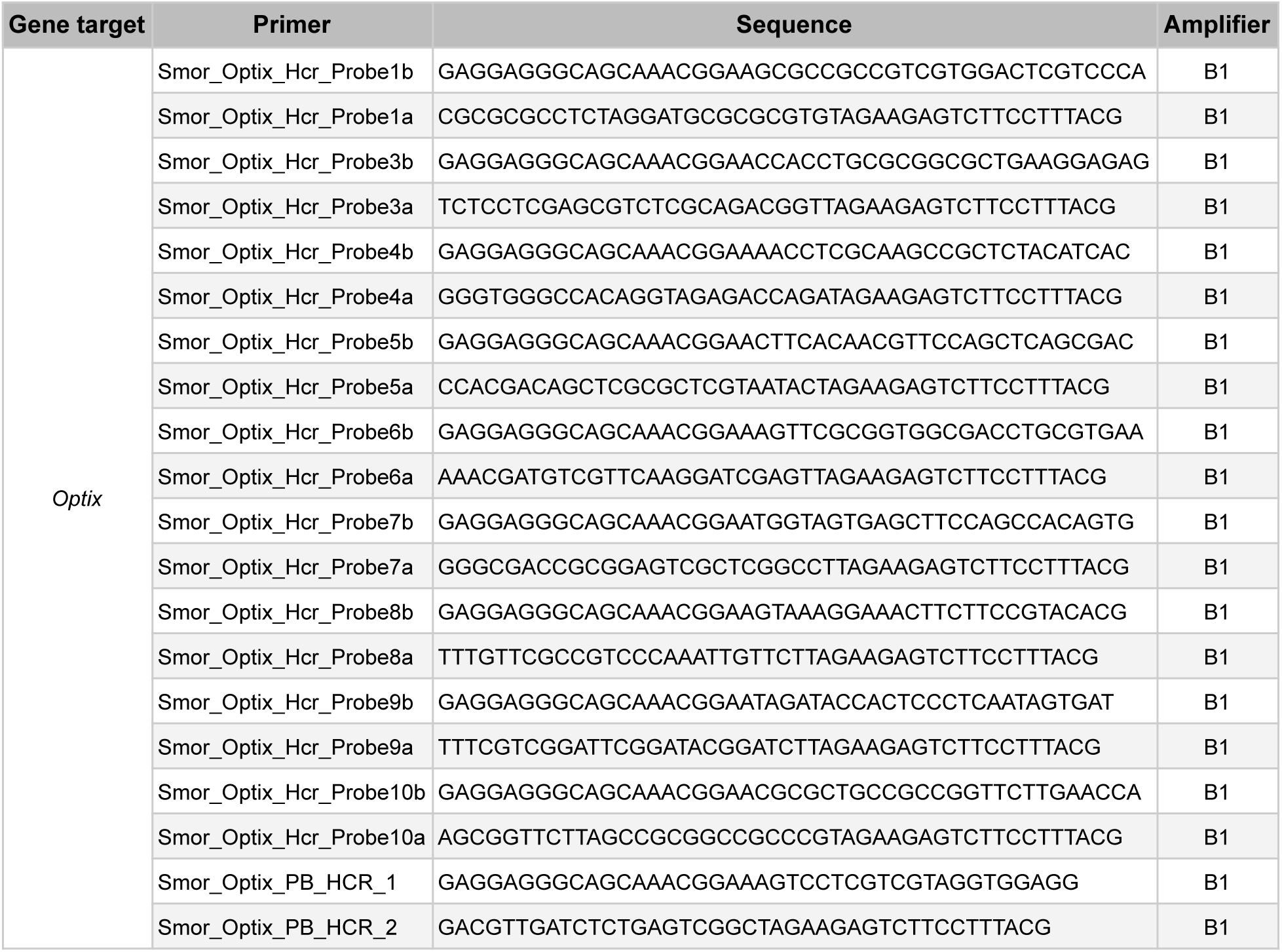
List of oligonucleotide used for the generation of *optix* HCR probes.

## References

1. Scott, J.A. (1992). The butterflies of North America: a natural history and field guide (Stanford University Press).

2. Morris, J., Hanly, J.J., Martin, S.H., Van Belleghem, S.M., Salazar, C., Jiggins, C.D., and Dasmahapatra, K.K. (2020). Deep convergence, shared ancestry, and evolutionary novelty in the genetic architecture of Heliconius mimicry. Genetics 216, 765–780. 10.1534/genetics.120.303611.

3. Reed, R.D., Papa, R., Martin, A., Hines, H.M., Counterman, B.A., Pardo-Diaz, C., Jiggins, C.D., Chamberlain, N.L., Kronforst, M.R., Chen, R., et al. (2011). optix Drives the Repeated Convergent Evolution of Butterfly Wing Pattern Mimicry. Science 333, 1137–1141. 10.1126/science.1208227.

4. Pardo-Diaz, C., Salazar, C., Baxter, S.W., Merot, C., Figueiredo-Ready, W., Joron, M., McMillan, W.O., and Jiggins, C.D. (2012). Adaptive Introgression across Species Boundaries in Heliconius Butterflies. PLOS Genetics 8, e1002752. 10.1371/journal.pgen.1002752.

5. Heliconius Genome Consortium (2012). Butterfly genome reveals promiscuous exchange of mimicry adaptations among species. Nature 487, 94–98.

6. Van Belleghem, S.M., Rastas, P., Papanicolaou, A., Martin, S.H., Arias, C.F., Supple, M.A., Hanly, J.J., Mallet, J., Lewis, J.J., Hines, H.M., et al. (2017). Complex modular architecture around a simple toolkit of wing pattern genes. Nature Ecology & Evolution 1.

7. Wallbank, R.W.R., Baxter, S.W., Pardo-Diaz, C., Hanly, J.J., Martin, S.H., Mallet, J., Dasmahapatra, K.K., Salazar, C., Joron, M., Nadeau, N., et al. (2016). Evolutionary Novelty in a Butterfly Wing Pattern through Enhancer Shuffling. PLOS Biology 14, e1002353. 10.1371/journal.pbio.1002353.

8. Cicconardi, F., Milanetti, E., Pinheiro de Castro, E.C., Mazo-Vargas, A., Van Belleghem, S.M., Ruggieri, A.A., Rastas, P., Hanly, J., Evans, E., Jiggins, C.D., et al. (2023). Evolutionary dynamics of genome size and content during the adaptive radiation of Heliconiini butterflies. Nat Commun 14, 5620. 10.1038/s41467-023-41412-5.

9. Dunford, J.C. (2009). Taxonomic overview of the greater fritillary genus Speyeria Scudder and the atlantis-hesperis species complexes, with species accounts, type images, and relevant literature (Lepidoptera: Nymphalidae). Insecta Mundi 90, 1–74.

10. Ren, A., Day, C.R., Hanly, J.J., Counterman, B.A., Morehouse, N., and Martin, A. (2020). Convergent evolution of broadband reflectors underlies metallic coloration in butterflies. Frontiers in Ecology and Evolution 8, 206.

11. Prakash, A., Finet, C., Banerjee, T.D., Saranathan, V., and Monteiro, A. (2022). Antennapedia and optix regulate metallic silver wing scale development and cell shape in Bicyclus anynana butterflies. Cell Reports 40, 111052. 10.1016/j.celrep.2022.111052.

12. Simonsen, T.J. (2007). Comparative morphology and evolutionary aspects of the reflective under wing scale-pattern in Fritillary butterflies (Nymphalidae: Argynnini). Zoologischer Anzeiger-A Journal of Comparative Zoology 246, 1–10.

13. Roberts, W.M. (2000). Dimorphism for silvering in Mormon fritillary butterflies (Speyeria mormonia) boisduval (Lepidoptera: Nymphalidae: Argynnini).

14. Hammond, P.C., McCorkle, D.V., and Bergman, W. (2013). Hybridization studies of genomic compatibility and phenotypic expression in the greater fritillary butterflies (Nymphalidae: Argynnini). The Journal of the Lepidopterists’ Society 67, 263–273.

15. Hammond, P.C., McCorkle, D.V., and Bergman, W. (2020). Additional Hybridization Studies of Genomic Compatibility and Phenotypic Expression in the Genus Speyeria (Nymphalidae: Argynnini). The Journal of the Lepidopterists’ Society 74, 133–153.

16. Tsai, C.-C., Childers, R.A., Shi, N.N., Ren, C., Pelaez, J.N., Bernard, G.D., Pierce, N.E., and Yu, N. (2020). Physical and behavioral adaptations to prevent overheating of the living wings of butterflies. Nature communications 11, 1–14.

17. Zhang, L., Mazo-Vargas, A., and Reed, R.D. (2017). Single master regulatory gene coordinates the evolution and development of butterfly color and iridescence. Proceedings of the National Academy of Sciences 114, 10707–10712.

18. Orteu, A., Hornett, E.A., Reynolds, L.A., Kemp, D.J., Gloder, G., Warren, I.A., Hurst, G.D.D., Martin, S.H., and Jiggins, C.D. (2024). Optix and cortex/ivory/mir-193 again: the repeated use of two mimicry hotspot loci. Proceedings of the Royal Society B: Biological Sciences 291, 20240627. 10.1098/rspb.2024.0627.

19. Martin, A., McCulloch, K.J., Patel, N.H., Briscoe, A.D., Gilbert, L.E., and Reed, R.D. (2014). Multiple recent co-options of Optix associated with novel traits in adaptive butterfly wing radiations. EvoDevo 5, 1–14. 10.1186/2041-9139-5-7.

20. Boggs, C.L. (1987). Demography of the unsilvered morph of Speyeria mormonia in Colorado, USA. Journal of The Lepidopterists’ Society 41, 94–97.

21. Ando, T., and Fujiwara, H. (2013). Electroporation-mediated somatic transgenesis for rapid functional analysis in insects. Development 140, 454–458. 10.1242/dev.085241.

22. Pavlidis, P., Živković, D., Stamatakis, A., and Alachiotis, N. (2013). SweeD: Likelihood-Based Detection of Selective Sweeps in Thousands of Genomes. Molecular Biology and Evolution 30, 2224–2234. 10.1093/molbev/mst112.

23. Przeworski, M. (2002). The Signature of Positive Selection at Randomly Chosen Loci. Genetics 160, 1179–1189. 10.1093/genetics/160.3.1179.

24. Moest, M., Van Belleghem, S.M., James, J.E., Salazar, C., Martin, S.H., Barker, S.L., Moreira, G.R., Mérot, C., Joron, M., Nadeau, N.J., et al. (2020). Selective sweeps on novel and introgressed variation shape mimicry loci in a butterfly adaptive radiation. PLoS Biology 18, e3000597.

25. Martin, S.H., and Belleghem, S.M.V. (2017). Exploring Evolutionary Relationships Across the Genome Using Topology Weighting. Genetics 206, 429–438. 10.1534/genetics.116.194720.

26. Martin, S.H., Davey, J.W., and Jiggins, C.D. (2015). Evaluating the Use of ABBA–BABA Statistics to Locate Introgressed Loci. Molecular Biology and Evolution 32, 244–257. 10.1093/molbev/msu269.

27. Zhang, J., Cong, Q., Shen, J., Opler, P.A., and Grishin, N.V. (2020). Genomic evidence suggests further changes of butterfly names. The taxonomic report of the International Lepidoptera Survey 8, 7.

28. Blount, Z.D., Lenski, R.E., and Losos, J.B. (2018). Contingency and determinism in evolution: Replaying life’s tape. Science 362, eaam5979. 10.1126/science.aam5979.

29. Stern, D.L., and Orgogozo, V. (2009). Is genetic evolution predictable? Science 323, 746–751.

30. Bohutínská, M., and Peichel, C.L. (2024). Divergence time shapes gene reuse during repeated adaptation. Trends in Ecology & Evolution 39, 396–407. 10.1016/j.tree.2023.11.007.

31. Martin, A., and Orgogozo, V. (2013). The loci of repeated evolution: a catalog of genetic hotspots of phenotypic variation. Evolution 67, 1235–1250.

32. Van Belleghem, S.M., Ruggieri, A.A., Concha, C., Livraghi, L., Hebberecht, L., Rivera, E.S., Ogilvie, J.G., Hanly, J.J., Warren, I.A., Planas, S., et al. (2023). High level of novelty under the hood of convergent evolution. Science 379, 1043–1049. 10.1126/science.ade0004.

33. Lewis, J.J., Geltman, R.C., Pollak, P.C., Rondem, K.E., Van Belleghem, S.M., Hubisz, M.J., Munn, P.R., Zhang, L., Benson, C., Mazo-Vargas, A., et al. (2019). Parallel evolution of ancient, pleiotropic enhancers underlies butterfly wing pattern mimicry. Proc. Natl. Acad. Sci. U.S.A. 116, 24174–24183. 10.1073/pnas.1907068116.

34. Huber, B., Whibley, A., Poul, Y.L., Navarro, N., Martin, A., Baxter, S., Shah, A., Gilles, B., Wirth, T., McMillan, W.O., et al. (2015). Conservatism and novelty in the genetic architecture of adaptation in Heliconius butterflies. Heredity 114, 515–524.

35. Kumar, S., Suleski, M., Craig, J.M., Kasprowicz, A.E., Sanderford, M., Li, M., Stecher, G., and Hedges, S.B. (2022). TimeTree 5: an expanded resource for species divergence times. Molecular biology and evolution 39, msac174.

36. Zhang, W., Dasmahapatra, K.K., Mallet, J., Moreira, G.R., and Kronforst, M.R. (2016). Genome-wide introgression among distantly related Heliconius butterfly species. Genome biology 17, 1.

37. Edelman, N.B., Frandsen, P.B., Miyagi, M., Clavijo, B., Davey, J., Dikow, R.B., García-Accinelli, G., Belleghem, S.M.V., Patterson, N., Neafsey, D.E., et al. (2019). Genomic architecture and introgression shape a butterfly radiation. Science 366, 594–599. 10.1126/science.aaw2090.

39. Polic, D., Yıldırım, Y., Merilaita, S., Franzén, M., and Forsman, A. (2024). Genetic structure, UV-vision, wing coloration and size coincide with colour polymorphism in Fabriciana adippe butterflies. Molecular Ecology 33, e17272. 10.1111/mec.17272.

40. Banerjee, T.D., Finet, C., Seah, K.S., and Monteiro, A. (2024). Optix regulates nanomorphology of butterfly scales primarily via its effects on pigmentation. Front. Ecol. Evol. 12. 10.3389/fevo.2024.1392050.

41. Zhang, L., Mazo-Vargas, A., and Reed, R.D. (2017). Single master regulatory gene coordinates the evolution and development of butterfly color and iridescence. PNAS 114, 10707–10712. 10.1073/pnas.1709058114.

42. Thayer, R.C., Allen, F.I., and Patel, N.H. (2020). Structural color in Junonia butterflies evolves by tuning scale lamina thickness. Elife 9, e52187.

43. Schwanwitsch, B.N. (1943). Stereomorphism in Cryptic Coloration of Lepidoptera. Nature 152, 508–508. 10.1038/152508a0.

44. Kjernsmo, K., Whitney, H.M., Scott-Samuel, N.E., Hall, J.R., Knowles, H., Talas, L., and Cuthill, I.C. (2020). Iridescence as Camouflage. Current Biology 30, 551–555.e3. 10.1016/j.cub.2019.12.013.

45. Kelley, J.L., Kelley, L.A., and Badcock, D.R. (2022). 3D animal camouflage. Trends in Ecology & Evolution 37, 628–631. 10.1016/j.tree.2022.04.001.

46. Henríquez-Piskulich, P., Stuart-Fox, D., Elgar, M., Marusic, I., and Franklin, A.M. (2023). Dazzled by shine: gloss as an antipredator strategy in fast moving prey. Behavioral Ecology 34, 862–871. 10.1093/beheco/arad046.

47. Kjernsmo, K., Lim, A.M., Middleton, R., Hall, J.R., Costello, L.M., Whitney, H.M., Scott-Samuel, N.E., and Cuthill, I.C. (2022). Beetle iridescence induces an avoidance response in naïve avian predators. Animal Behaviour 188, 45–50. 10.1016/j.anbehav.2022.04.005.

48. Zhang, J., Cong, Q., Shen, J., Song, L., Gott, R.J., Boyer, P., Guppy, C.S., Kohler, S., Lamas, G., Opler, P.A., et al. (2022). Taxonomic discoveries enabled by genomic analysis of butterflies. The taxonomic report of the International Lepidoptera Survey 10, 1. 10.5281/zenodo.7160429.

49. Fick, S.E., and Hijmans, R.J. (2017). WorldClim 2: new 1-km spatial resolution climate surfaces for global land areas. International Journal of Climatology 37, 4302–4315. 10.1002/joc.5086.

50. Aiello-Lammens, M.E., Boria, R.A., Radosavljevic, A., Vilela, B., and Anderson, R.P. (2015). spThin: an R package for spatial thinning of species occurrence records for use in ecological niche models. Ecography 38, 541–545. 10.1111/ecog.01132.

51. Kass, J.M., Muscarella, R., Galante, P.J., Bohl, C.L., Pinilla-Buitrago, G.E., Boria, R.A., Soley-Guardia, M., and Anderson, R.P. (2021). ENMeval 2.0: Redesigned for customizable and reproducible modeling of species’ niches and distributions. Methods in Ecology and Evolution 12, 1602–1608. 10.1111/2041-210X.13628.

51. Jarvis, A., Guevara, E., Reuter, H.I., and Nelson, A.D. (2008). Hole-filled SRTM for the globe: version 4: data grid. Web publication/site, CGIAR Consortium for Spatial Information. http://srtm.csi.cgiar.org/ http://srtm.csi.cgiar.org/.

53. Hersbach, H., Bell, B., Berrisford, P., Biavati, G., Horányi, A., Muñoz Sabater, A., Nicolas, J., Peubey, C., Radu, R., Rozum, I., et al. ERA5 monthly averaged data on single levels from 1940 to present. Copernicus Climate Change Service (C3S) Climate Data Store (CDS). 10.24381/cds.f17050d7.

54. Mattoon, S.O., Davis, R.D., and Spencer, O.D. (1971). Rearing techniques for species of Speyeria (Nymphalidae). Journal of the Lepidopterists’ Society 25, 247–255.

55. Pedersen, B.S., and Quinlan, A.R. (2018). Mosdepth: quick coverage calculation for genomes and exomes. Bioinformatics 34, 867–868.

56. Li, H. (2013). Aligning sequence reads, clone sequences and assembly contigs with BWA-MEM. Preprint at arXiv, 10.48550/arXiv.1303.3997 10.48550/arXiv.1303.3997.

56. Broad Institute, G.R. (2019). Picard Toolkit. https://broadinstitute.github.io/picard/ https://broadinstitute.github.io/picard/.

57. Van der Auwera, G.A., Carneiro, M.O., Hartl, C., Poplin, R., del Angel, G., Levy-Moonshine, A., Jordan, T., Shakir, K., Roazen, D., Thibault, J., et al. (2013). From FastQ Data to High-Confidence Variant Calls: The Genome Analysis Toolkit Best Practices Pipeline. Current Protocols in Bioinformatics 43, 11.10.1–11.10.33. 10.1002/0471250953.bi1110s43.

59. Danecek, P., Bonfield, J.K., Liddle, J., Marshall, J., Ohan, V., Pollard, M.O., Whitwham, A., Keane, T., McCarthy, S.A., Davies, R.M., et al. (2021). Twelve years of SAMtools and BCFtools. GigaScience 10, giab008. 10.1093/gigascience/giab008.

60. Browning, B.L., Zhou, Y., and Browning, S.R. (2018). A One-Penny Imputed Genome from Next-Generation Reference Panels. The American Journal of Human Genetics 103, 338–348. 10.1016/j.ajhg.2018.07.015.

61. Browning, B.L., Tian, X., Zhou, Y., and Browning, S.R. (2021). Fast two-stage phasing of large-scale sequence data. The American Journal of Human Genetics 108, 1880–1890. 10.1016/j.ajhg.2021.08.005.

62. Chang, C.C., Chow, C.C., Tellier, L.C., Vattikuti, S., Purcell, S.M., and Lee, J.J. (2015). Second-generation PLINK: rising to the challenge of larger and richer datasets. GigaScience 4, s13742–015-0047–0048. 10.1186/s13742-015-0047-8.

63. Zhou, X., and Stephens, M. (2012). Genome-wide efficient mixed-model analysis for association studies. Nat Genet 44, 821–824. 10.1038/ng.2310.

64. Whiting, J. (2022). Genotype Plot. Version v0.2.1. 10.5281/zenodo.5913504 10.5281/zenodo.5913504.

65. Cheng, H., Concepcion, G.T., Feng, X., Zhang, H., and Li, H. (2021). Haplotype-resolved de novo assembly using phased assembly graphs with hifiasm. Nat Methods 18, 170–175. 10.1038/s41592-020-01056-5.

65. Heryanto, C., Hanly, J.J., Mazo-Vargas, A., Tendolkar, A., and Martin, A. (2022). Mapping and CRISPR homology-directed repair of a recessive white eye mutation in Plodia moths. iScience, 103885.

67. Choi, H.M., Schwarzkopf, M., Fornace, M.E., Acharya, A., Artavanis, G., Stegmaier, J., Cunha, A., and Pierce, N.A. (2018). Third-generation in situ hybridization chain reaction: multiplexed, quantitative, sensitive, versatile, robust. Development 145, dev165753.

68. Kuehn, E., Clausen, D.S., Null, R.W., Metzger, B.M., Willis, A.D., and Özpolat, B.D. (2022). Segment number threshold determines juvenile onset of germline cluster expansion in Platynereis dumerilii. Journal of Experimental Zoology Part B: Molecular and Developmental Evolution 338, 225–240. 10.1002/jez.b.23100.

69. Bruce, H.S., Jerz, G., Kelly, S., McCarthy, J., Pomerantz, A., Senevirathne, G., Sherrard, A., Sun, D.A., Wolff, C., and Patel, N.H. (2021). Hybridization chain reaction (HCR) in situ protocol. protocols. io. 10.17504/protocols.io.bunznvf6.

70. Fujiwara, H., and Nishikawa, H. (2016). Functional analysis of genes involved in color pattern formation in Lepidoptera. Current opinion in insect science 17, 16–23.

